# Interpretable deep learning architectures for improving drug response prediction performance: myth or reality?

**DOI:** 10.1101/2022.10.03.510614

**Authors:** Yihui Li, David Earl Hostallero, Amin Emad

## Abstract

Motivation: Recent advances in deep learning model development have enabled more accurate prediction of drug response in cancer. However, the black-box nature of these models still remains a hurdle in their adoption for precision cancer medicine. Recent efforts have focused on making these models interpretable by incorporating signaling pathway information in model architecture. While these models improve interpretability, it is unclear whether this higher interpretability comes at the cost of less accurate predictions, or a prediction improvement can also be obtained. Results: In this study, we comprehensively and systematically assessed four state-of-the-art interpretable models developed for drug response prediction to answer this question using three pathway collections. Our results showed that models that explicitly incorporate pathway information in the form of a latent layer perform worse compared to models that incorporate this information implicitly. Moreover, in most evaluation setups the best performance is achieved using a simple black-box model. In addition, replacing the signaling pathways with randomly generated pathways shows a comparable performance for the majority of these interpretable models. Our results suggest that new interpretable models are necessary to improve the drug response prediction performance. In addition, the current study provides different baseline models and evaluation setups necessary for such new models to demonstrate their superior prediction performance. Availability and Implementation: Implementation of all methods are provided in https://github.com/Emad-COMBINE-lab/InterpretableAI_for_DRP. Generated uniform datasets are in https://zenodo.org/record/7101665#.YzS79HbMKUk. Contact

Supplementary Information: Online-only supplementary data is available at the journal’s website.

## Introduction

Machine learning models have found various applications in medicine, including drug repositioning (Jarada, et al., 2020), drug discovery (Vamathevan, et al., 2019), gene prioritization (Emad, et al., 2017; Zhang, et al., 2021), and drug response prediction (Adam, et al., 2020; Ballester, et al., 2022; Costello, et al., 2014; Huang, et al., 2020). Models for drug response prediction (DRP) are typically trained using various data modalities such as molecular ‘omics’ profiles of samples (e.g., cancer cell lines or tumors), drug representations, and network information (Adam, et al., 2020; Ballester, et al., 2022; Guvenc Paltun, et al., 2021). In recent years, various models have been proposed using deep learning (DL) for drug response prediction (Baptista, et al., 2021; Chen and Zhang, 2022; El Khili, et al., 2022; Hostallero, et al., 2022; Hostallero, et al., 2021). In spite of their success in their perspective tasks, most DL models are considered as “black-boxes” with inner operations that are difficult to interpret. This characteristic of DL models is undesirable for applications in the biomedical field, as identifying the set of biological features that contribute to the model prediction outputs and understanding the relationship between these features are crucial when conducting further experimental studies to validate these computational findings. To address these challenges, the concept of interpretable artificial intelligence (Azodi, et al., 2020; Barredo Arrieta, et al., 2020; Malioutov, et al., 2017) has been introduced to create models that can achieve both high performance and interpretability.

In the context of DRP, model interpretability can be achieved in two ways: 1) using post-hoc analysis to determine feature attributions and identify important features without explicitly incorporating prior knowledge in model architecture, and 2) integrating prior knowledge (e.g., signaling pathways) to add meaningful structure to the model, which can then be interpreted (for example using post-hoc feature importance methods). While we and others have successfully used the former strategy in DRP (Hostallero, et al., 2022; Hostallero, et al., 2021) and other applications (Caruana, et al., 2015; Che, et al., 2016), the latter strategy can potentially allow the interpretability to go one step further to provide systems biology insights regarding the mechanisms involved in response to drug treatments. Incorporating prior information such as biological pathway and subsystem information allows the model embeddings to reflect subsystem activities and state changes, which can then be computationally or experimentally investigated to reveal different biological mechanisms that confer specific drug sensitivities (Kuenzi, et al., 2020). In fact, post-hoc feature importance analysis can be incorporated in these models to identify not only important input features, but also embeddings that reflect crucial subsystems for cellular response to a particular drug.

The models that incorporate pathway information have generated valuable insights regarding drugs’ mechanisms of action and gene-pathway relationships, some of which have been validated experimentally (Kuenzi, et al., 2020). However, there have been conflicting reports on their ability in providing accurate drug response predictions (Deng, et al., 2020; Jin and Nam, 2021; Kuenzi, et al., 2020; Snow, et al., 2021; Tang and Gottlieb, 2021; Zhang, et al., 2021). Ideally, interpretability should not come at the expense of prediction performance, since a lower prediction performance of interpretable models may reflect that the black-box models are better capable at extracting patterns of the data and incorporating informative signals that are not being utilized by the more interpretable models. For example, consider a hypothetical model that is completely interpretable, but generates random drug response predictions that do not reflect the measured drug responses of samples. No matter how interpretable this model may be, the insights obtained from it is not going to reflect the biological and chemical mechanisms involved in drug response.

Recognizing the intertwined relationship between interpretability and performance, the majority of recent models that incorporate pathway information for better interpretability have also sought and reported an improved prediction performance (Deng, et al., 2020; Jin and Nam, 2021; Snow, et al., 2021; Tang and Gottlieb, 2021; Zhang, et al., 2021). On the other hand, some studies have reported comparable or slightly worse model performance after incorporating pathway information (Kuenzi, et al., 2020). However, it is rather difficult to gauge the (potential) contribution of pathway information in DRP performance from the original studies, due to differences between data used in each study, their evaluation setup, and in many cases a lack of appropriate baseline models to act as control. To investigate these inconsistent findings in state-of-the-art models, we conducted a study that comprehensively evaluates the effect of pathway incorporation on performance of DRP models and aims to answer five main questions:

1. Does the inclusion of biological pathway information improve model performance when evaluated strictly and comprehensively?
2. Which type of pathway incorporation strategy is best capable of improving the performance?
3. Are interpretable models better suited for prediction of response of unseen cell lines or unseen drugs?
4. Can the performance of the interpretable models be attributed to biological information present in the pathway datasets, or a similar improvement can be also achieved through the use of randomly generated pathways, reflecting a technical (instead of a biological) origin for the performance?
5. What pathway database is most helpful in improving model performance?

To answer the proposed questions, we performed 189 experiments evaluating 21 computational models with three pathway collections (Kanehisa and Goto, 2000), (Schaefer, et al., 2009), (Fabregat, et al., 2017) and under three data splitting strategies. The models included four state-of-the-art interpretable DL architectures that incorporate pathway information (Deng, et al., 2020; Jin and Nam, 2021; Tang and Gottlieb, 2021; Zhang, et al., 2021) (henceforth pathway-based models) and four variants of them, as well as thirteen baseline models that can evaluate the performance of these models from different angles (discussed in Methods). We selected these interpretable models since they use similar type of information for cancer cell lines (CCLs) and drugs and utilize gene-pathway membership in their architectures, allowing us to compare them fairly and comprehensively. Moreover, they represent two categories of strategies to incorporate pathway information in DL architectures: methods that use a pathway layer connecting genes to pathway nodes (*explicit* models) such as PathDNN (Deng, et al., 2020) and ConsDeepSignaling (CDS) (Zhang, et al., 2021), and those that do not directly use a pathway layer (*implicit* models) such as HiDRA (Jin and Nam, 2021), and PathDSP (Tang and Gottlieb, 2021).

Our baseline models included a traditional machine learning model (random forests), a black-box fully connected neural network with a similar architecture to those of the interpretable models, as well as “naive” predictors and “random-pathway” predictors, two important baselines that have been largely overlooked in previous studies. The naive predictor uses the average drug response of samples in the training set and reports that for each testing sample. This baseline is particularly important in controlling for inflation of prediction performance due to distinct range of log IC50 (natural log of the half maximal inhibitory concentration, a drug sensitivity measure) of different drugs. In other words, it is possible to obtain a good approximation of drug response by simply knowing the identity of the drug, resulting in artificially inflated performance metrics. Each random-pathway predictor exactly matches the architecture and pipeline of an interpretable model, but randomly assigns genes to pathways, while preserving the size of each pathway. These baselines allow us to determine whether potential performance improvement of an interpretable model is truly due to the added value of the biological information, or instead is a technical artifact of modifying the model architecture.

Our analysis showed that overall, incorporating pathway information *does not* lead to improved prediction performance, confirming the observations reported by Kuenzi et al. (Kuenzi, et al., 2020) for their proposed model. In particular, in many cases a simple black-box multilayer perceptron (MLP) achieves the best performance. Moreover, even in instances that performance improvement compared to an MLP or a naive predictor was observed, a similar performance was achieved using randomly generated pathways. This suggests that such improvements should not be attributed to the biological information carried by pathway collections and is likely a technical artifact. We also observed that the strategy used to include pathway information in the models has a significant influence on the performance, and explicit models seem to perform worse compared to implicit models. Finally, Reactome pathways seemed to provide slightly better predictions compared to other pathway collections.

## Methods

### Data preprocessing and uniform dataset formation

To form uniform datasets for our analyses, we first evaluated different data modalities and datasets used by each of the pathway-based models (Supplementary Table S1). In these studies, gene expression (GEx), somatic mutation (Mut), and copy number variation (CNV) of samples were used, while for drugs their targets (T) or their Morgan fingerprints (FP) capturing their chemical structure were used. In order to maintain fairness and consistency of model performance comparisons, for each choice of pathway collection we compiled a uniform dataset that was used by all models evaluated in this study (three uniform datasets in total). These datasets are freely available in https://zenodo.org/record/7101665#.YzS79HbMKUk.

**Table 1:**
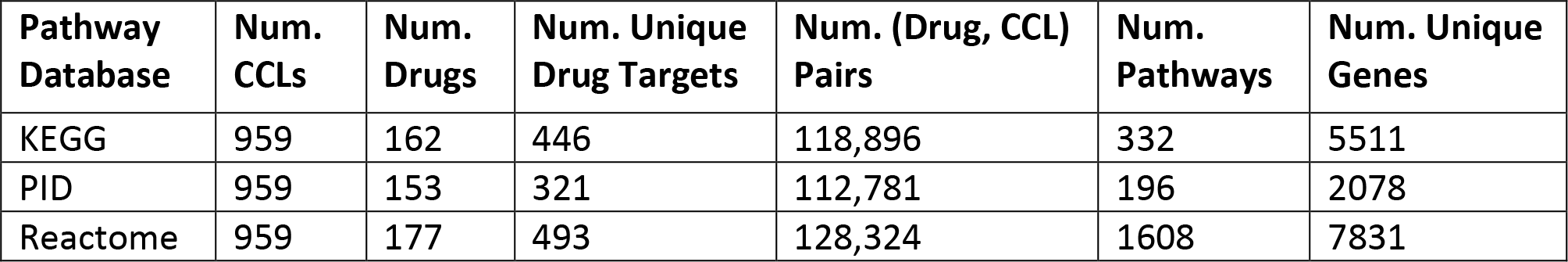
Summary of pathway-specific uniform datasets.

| Pathway Database | Num. CCLs | Num. Drugs | Num. Unique Drug Targets | Num. (Drug, CCL) Pairs | Num. Pathways | Num. Unique Genes |
| --- | --- | --- | --- | --- | --- | --- |
| KEGG | 959 | 162 | 446 | 118,896 | 332 | 5511 |
| PID | 959 | 153 | 321 | 112,781 | 196 | 2078 |
| Reactome | 959 | 177 | 493 | 128,324 | 1608 | 7831 |

We collected GEx, Mut, CNV, and drug sensitivity data (in the form of log IC50) of 959 cancer cell lines (CCLs) from Genomics of Drug Sensitivity in Cancer (GDSC) (Yang, et al., 2013) database. We obtained drug target information from STITCH (Szklarczyk, et al., 2016) and drug structural data from PubChem (Kim, et al., 2021). Protein-protein interactions (PPI) that were used by one of the models were obtained from the STRING database (Szklarczyk, et al., 2019) (only experimental

PPIs were used). Finally, gene-pathway membership information was obtained from KEGG (Kyoto Encyclopedia of Genes and Genomes) (Kanehisa and Goto, 2000), PID (Pathway Interaction Database) (Schaefer, et al., 2009), and Reactome (Fabregat, et al., 2017). Supplementary Table S2 outlines the data used in this study and their sources. We obtained drug response data in the form of log IC50 values and removed duplicate drugs that came from different experimental batches. In such cases, we kept the drug whose response was measured across a larger number of CCLs. We collected drug InChI (International Chemical Identifier) strings (Heller, et al., 2015) from PubChem and used the RDKit (Landrum, 2006) software to generate 512-bit Morgan fingerprints for these drugs. We obtained drug target data from the STITCH database, where we only kept drug targets with confidence score larger than 800 (out of 1000) and coming from the “experimental” and “database” channels.

**Table 2:**
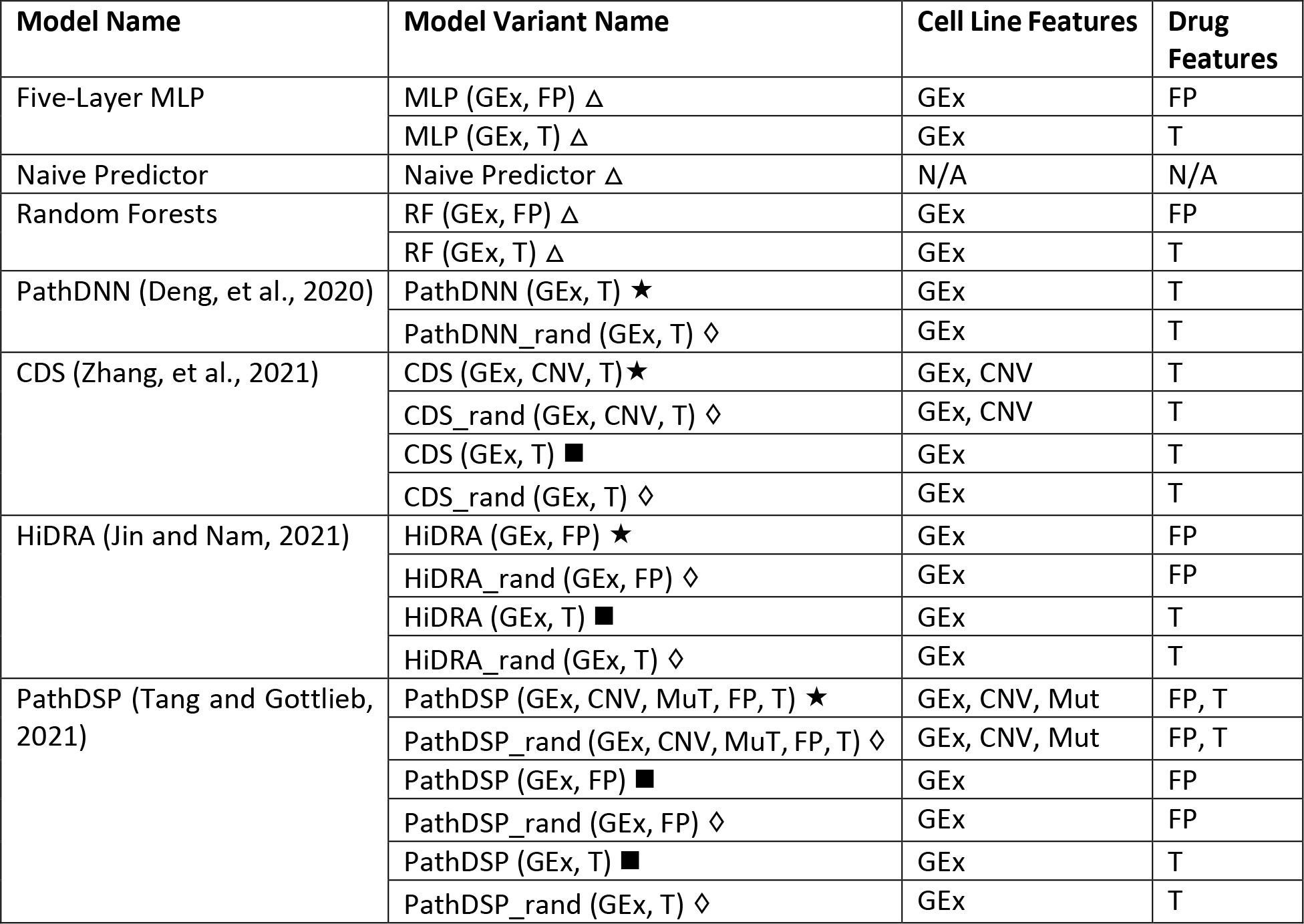
List of evaluated models. △ = universal baseline, ♢ = model-specific random pathway baseline, * = original pathway-based model, ▪ = model variant, GEx = gene expression, CNV = copy number variation, Mut = somatic mutation, T = drug target data, FP = Morgan fingerprint (drug structural data)

We performed log2(FPKM+1) normalization on the GEx data and removed genes whose expression showed low variability across different CCLs (standard deviation < 0.1). We also removed genes for which there were no somatic mutations, CNV, pathway information, drug target data, and STRING Experimental PPI information. This formed our common gene set (Figure 1A and 1B). In parallel, drug targets that were not present in the common gene set above or in the PPI network were excluded. Only drugs that had both log IC50 measurements and drug targets were kept in the final uniform datasets (Figure 1C and 1D). The PPI network was involved in the data preprocessing step as PathDSP (Tang and Gottlieb, 2021) incorporated it to perform pathway enrichment analysis. Since we needed the uniform datasets to be usable by all models, we included this step in the pre-processing procedure.

**Figure 1:**
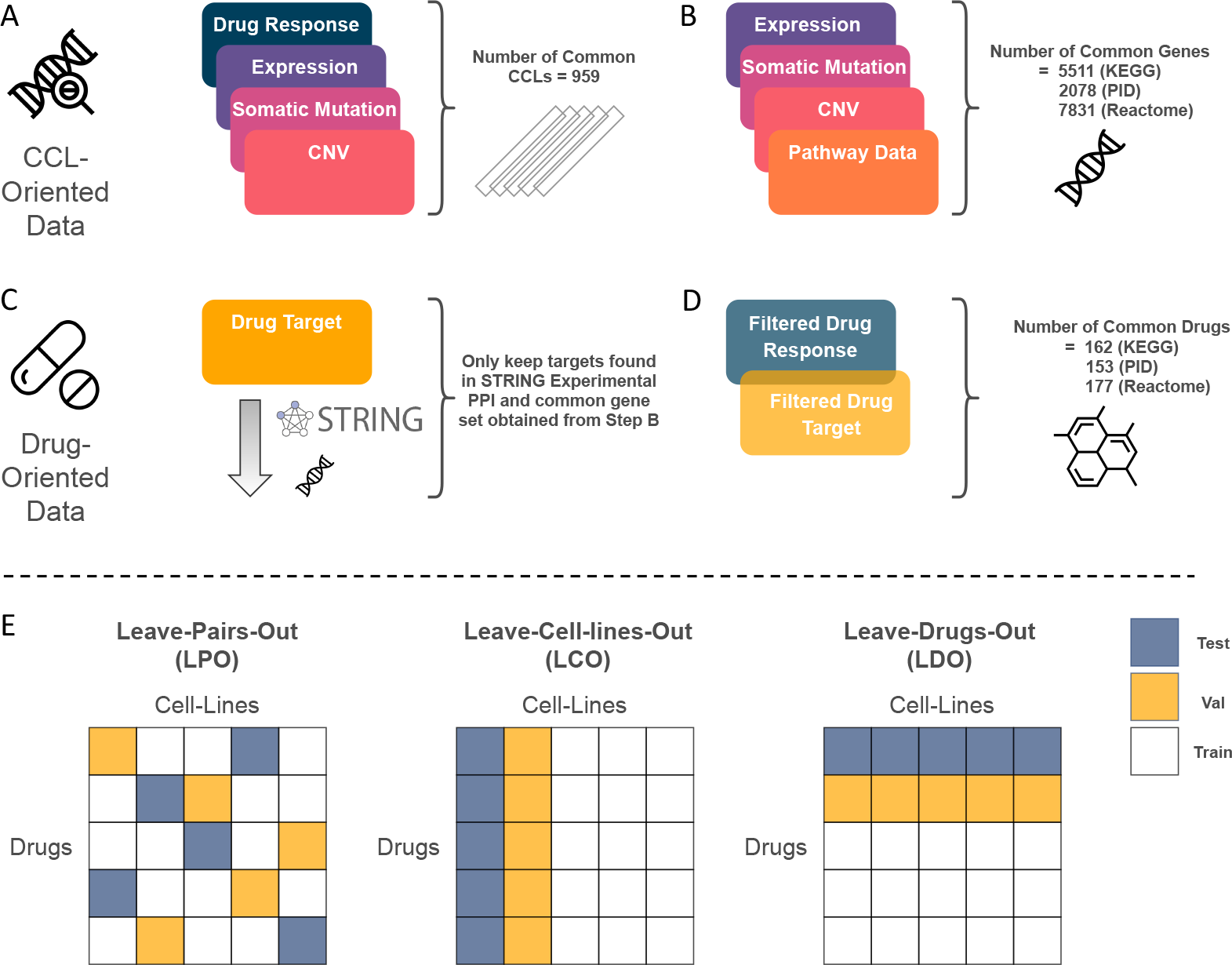
Construction of pathway-specific uniform datasets and data splitting approaches. (A) cancer cell lines (CCLs) with available data for drug response, gene expression, somatic mutation, and copy number variation (CNV) were selected. (B) Genes shared between different sources of data were identified. Genes that were not present in any pathway were removed. (C) Drug target genes that were not found in the common gene set obtained from Step B and the STRING Experimental protein-protein interaction (PPI) network were removed. (D) Drugs and small molecules that had measured log IC50 values and drug target information were selected. E) Model input data was split into five folds, with the training, validation, and test set ratio of 3: 1: 1. Folds in the leave-pairs-out (LPO) validation scheme are formed by randomly selecting mutually exclusive (CCL, drug) pairs, whereas in leave-cell-lines-out (LCO) and leave-drugs-out (LDO) validation schemes, mutually exclusive cell lines and drugs are randomly selected, respectively.

Figure 1 illustrates the process of constructing the pathway-specific uniform datasets. Since each source of pathway collection contained different number of genes, the final dataset for each collection was slightly different. Table 1 summarizes the number of CCLs, drugs, genes, and pathways for each pathway collection in the uniform dataset, while Supplementary Table S3 provides details about CCLs and drugs.

**Table 3:** Performance of pathway-based models using KEGG collection, with gene expression (GEx) and drug targets (T) as inputs. The mean and standard deviations (std) are calculated across cancer cell lines (CCLs). The best performing model is bold-faced, while worst performing model is underlined. Models are ranked by their leave-cell-lines-out (LCO) RMSE. The following symbols are used in this table: △ = universal baseline, * = original pathway-based model, ▪ = model variant, ↑ = higher value indicates better performance, ↓ = lower value indicates better performance. ∗ The leave-drugs-out (LDO) Spearman’s correlation coefficient (SCC) cannot be calculated for the naive predictor since in this case it outputs the same value for all CCLs. For performance of these models based on Pearson correlation coefficient, R-squared, mean squared error (MSE), and concordance index, see Supplementary Table S4. Supplementary Figure S2 provides visualization of these values in the form of bar plots.

| Model Name | LCO |  | LDO |  | LPO |  |
| --- | --- | --- | --- | --- | --- | --- |
| | SCC ↑<br>(±std) | RMSE ↓<br>(±std) | SCC ↑<br>(±std) | RMSE ↓<br>(±std) | SCC ↑<br>(±std) | RMSE ↓<br>(±std) |
| PathDSP (GEx, T) ■ | **0.882**<br>**(±0.045)** | **1.283**<br>**(±0.230)** | 0.380<br>(±0.146) | **2.648**<br>**(±0.287)** | **0.876**<br>**(±0.074)** | **1.103**<br>**(±0.256)** |
| RF (GEx, T) $\Delta$ | 0.867<br>(±0.042) | 1.329<br>(±0.205) | <u>0.342</u><br>(±0.175) | 2.955<br>(±0.474) | 0.824<br>(±0.096) | 1.349<br>(±0.301) |
| HiDRA (GEx, T) ■ | 0.864<br>(±0.048) | 1.368<br>(±0.253) | 0.356<br>(±0.156) | <u>3.110</u><br>(±0.400) | 0.863<br>(±0.078) | 1.174<br>(±0.260) |
| Naive Predictor $\Delta$ | 0.871<br>(±0.045) | 1.373<br>(±0.292) | NA * | 2.826<br>(±0.274) | 0.855<br>(±0.087) | <u>1.742</u><br>(±0.280) |
| MLP (GEx, T) $\Delta$ | 0.858<br>(±0.042) | 1.420<br>(±0.231) | **0.382**<br>**(±0.160)** | 2.686<br>(±0.366) | 0.845<br>(±0.086) | 1.294<br>(±0.286) |
| PathDNN (GEx, T) $\star$ | 0.857<br>(±0.044) | 1.494<br>(±0.292) | <u>0.342</u><br>(±0.179) | 3.054<br>(±0.485) | 0.851<br>(±0.083) | 1.245<br>(±0.273) |
| CDS (GEx, T) ■ | <u>0.789</u><br>(±0.098) | <u>1.603</u><br>(±0.254) | 0.345<br>(±0.156) | 2.724<br>(±0.317) | <u>0.769</u><br>(±0.106) | 1.508<br>(±0.304) |

### Model evaluation and data split

We split our data randomly into five disjoint folds, where the training, validation, and test ratio was 3: 1: 1. The validation set was used for hyperparameter tuning and the test set was used for final model evaluation. The details of hyperparameter tuning, model training, and final architectures are provided in Supplementary File S2. We adopted three data splitting methods (validation schemes) to generate these folds: leave-pairs-out (LPO), leave-cell-lines-out (LCO), and leave-drugs-out (LDO), as depicted in Figure 1E. These three strategists were adopted to comprehensively assess the models for different drug response prediction tasks (for unseen (CCL, drug) pairs, unseen CCLs, and unseen drugs, respectively), and to determine in which one of these tasks (if any) pathway incorporation improves model prediction performance. To ensure fairness, same folds were used for all models.

We evaluated the performance of each model using two main performance measures: Spearman’s correlation coefficient (SCC) and root mean squared error (RMSE), but various other measures are also reported in supplementary tables. First, for a fixed CCL, the predicted values across all drugs of the test set were compared with the measured log IC50 values to calculate a CCL-specific performance measure (SCC or RMSE). Then, the mean and standard deviation of the performance measure was calculated across all CCLs.

### Overview of interpretable models and their variants

To study the effect of incorporating pathway information on drug response prediction, we selected four pathway-based state-of-the-art models: PathDNN (Deng, et al., 2020), ConsDeepSignaling (CDS) (Zhang, et al., 2021), HiDRA (Jin and Nam, 2021), and PathDSP (Tang and Gottlieb, 2021). We selected these models since 1) they use similar types of information for CCLs and drugs and utilize gene-pathway membership in their architectures (instead of other types of prior information such as hierarchical relationships of gene ontologies), 2) they all showed improved drug response prediction performance in their original studies compared to their black-box counterparts, and 3) they represent two important categories of implicit and explicit models (as discussed earlier). While other important models also exist (e.g., DrugCell (Kuenzi, et al., 2020)), they did not satisfy the conditions above. For example, DrugCell (unlike the models above) uses the hierarchical structure of gene ontologies and pathways, making it rather difficult to compare against the models above in a fair manner, since it takes advantage of more detailed information. Moreover, the original study of DrugCell showed that while including prior information improved interpretability of their model, it did not improve the performance of drug response prediction compared to its black-box counterpart. Due to the reasons above, we decided to exclude it from this analysis.

Models with an *explicit* pathway layer (e.g., PathDNN and CDS) typically define a gene and a pathway layer with connections between these layers reflecting gene-pathway membership (Figure 2A-2B). The input layer of this category of models contains drug and cell line features at gene level. As a result, only drug gene targets can be used with these models and Morgan Fingerprint (and other structural data) is not usable without altering the model architecture. The pathway layer is then connected to a group of fully connected layers to predict drug response for a given sample. The inclusion of the pathway layer allows identification of important pathways for a particular drug treatment or cancer type through post-hoc feature importance analysis.

**Figure 2:**
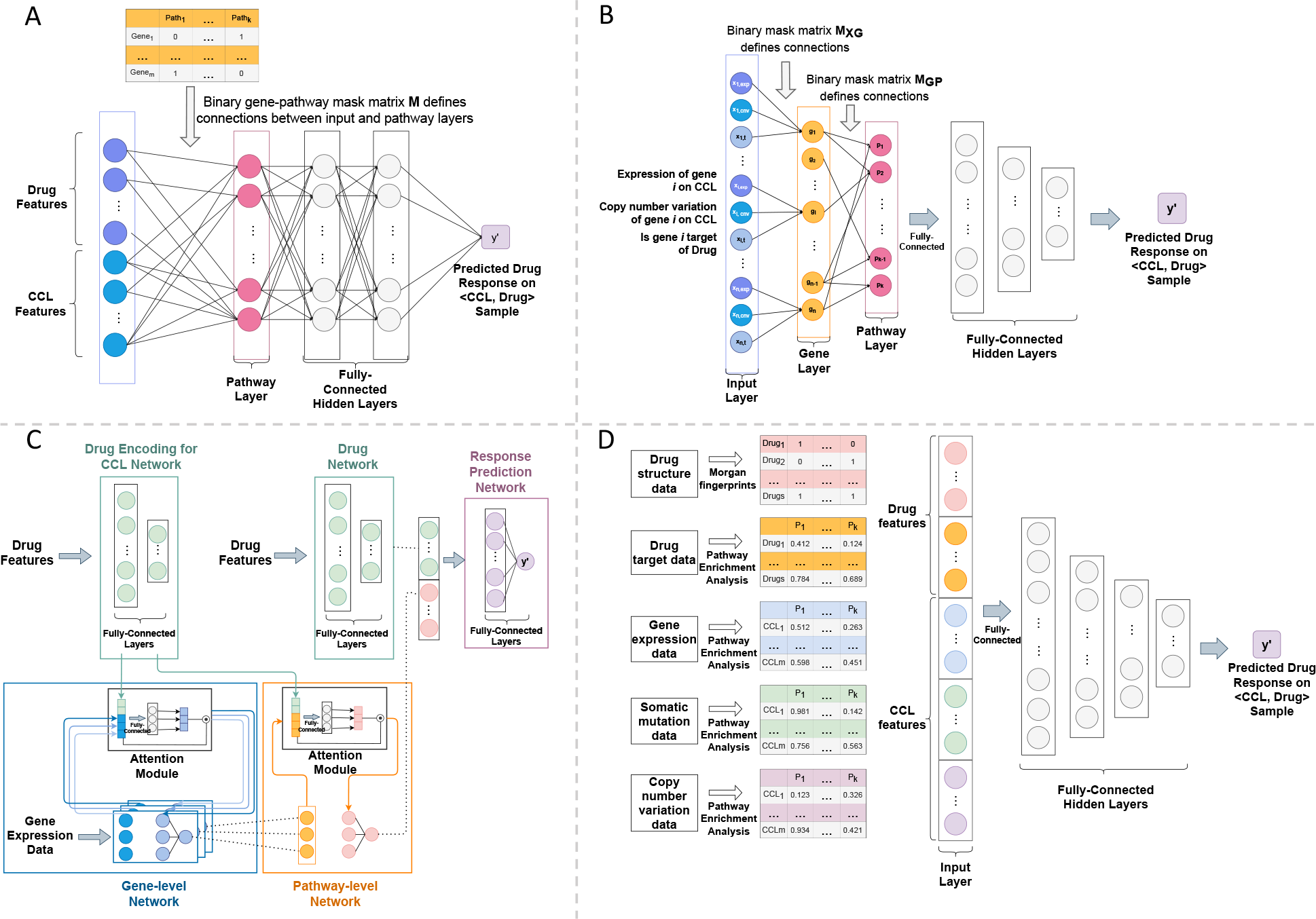
An overview of pathway-based models considered in this study. A) PathDNN uses cancer cell line (CCL) gene expression profiles as CCL features and drug target information as drug features. The input features (genes) are connected to the pathway nodes through gene-pathway membership. The pathway layer is followed by a set of fully connected layers. B) ConsDeepSignaling (CDS) takes gene expression profile and copy number variation as CCL features and drug target information as drug features. Each node in the gene layer represents a gene and is connected to its corresponding input features in the input layer (through connection matrix MXG). Connections between the gene and pathway layer are defined by gene-pathway membership (binary connection matrix MGP). A set of fully connected layers follow the pathway layer. C) HiDRA has a hierarchical network architecture. It uses gene expression profiles as CCL features. Drug target information and structural data can be both used as drug features. The pathway information is incorporated using an attention module, where a small neural network is dedicated to each pathway. Pathway activation scores are calculated by the gene-level network and are concatenated with drug feature embeddings learned by the drug encoding network to generate the final input to the drug response prediction network. D) In PathDSP, drug target, gene expression, somatic mutation, and copy number variation data are processed using pathway enrichment analysis to from matrices of enrichment scores, which act as input features to the model, which is a set of fully connected layers.

Models that *implicitly* incorporate pathway information take various forms. For example, HiDRA (Jin and Nam, 2021) uses a gene-level and pathway-level attention module to calculate pathway importance scores, where a small-scale neural network is dedicated to each pathway by only using features associated with the member genes of that specific pathway as inputs (Figure 2C). On the other hand, PathDSP uses a classic fully connected feedforward architecture, but the input features are pathway-enrichment scores rather than gene-level features (Figure 2D). See Supplementary Files S1 and S2 for details regarding models’ architectures and their training procedure.

Each of the pathway-based models used different data modalities in their original study (Supplementary Table S1). We tested all models using the three pathway collections discussed earlier. For implicit models, we tried both drug targets (T) and Morgan fingerprints (FP); however, for explicit models, only drug targets could be used due to their requirements that the drug features must be at gene level. For CCL features, we used all data modalities used by the original study. However, since gene expression data was used by all models (alone or in combination with other omics data, Supplementary Table S1), we also implemented model variants that only utilized GEx data. This ensured that one architecture is not given an unfair advantage due to access to a larger number of modalities. Table 2 provides a summary of all variations of the models considered in this study.

### Baseline Models

We used four types of baseline models to benchmark the pathway-based models and their variants. First, we used a multilayer perceptron (MLP) with five layers as a universal baseline for all models. This MLP represents a black-box feedforward neural network that is often used for benchmarking of other deep learning architectures (including the pathway-based models). Since all pathway-based models had a variant trained with GEx data, along with drug targets (or Morgan fingerprints), we trained two MLP models, MLP (GEx, FP) and MLP (GEx, T), representing the data input options above (Table 2).

The second type of baseline used in our study was a predictor that simply calculates the average drug sensitivity measure of samples in the training set and reports their average for all samples in the test set (henceforth referred as naive predictor). More specifically, the naive predictor does not use any CCL or drug features, but instead simply relies on the identity of the CCL or the drug (depending on the data splitting strategy). As shown in Supplementary Figure S1, in the LCO setup and for a (CCL, drug) pair in the test set, the naive predictor reports the average response of all CCLs in the training set to that drug. As a result, all CCLs in the test set will have the same response value for a drug (i.e., only the drug identity determines the response). On the other hand, in the LDO setup and for a (CCL, drug) pair, the average response of the CCL to all drugs in the training set is reported as the prediction (i.e., only the identity of the CCL determines the response). In the case of LPO, the averaging is done across all drugs and all CCLs corresponding to a (CCL, drug) in the test set. The naive predictors reveal the performance of a model that does not learn the relationship between the input features and output drug response and can control for inflation in the performance metrics.

The third type of baselines correspond to model-specific baselines that have the exact same architecture of a pathway-based model (with all their input data), but instead of gene-pathway membership information from pathway databases use randomly generated pathways. This type of baseline model (shown with a suffix of “_rand” in Table 2) allows us to determine if the (potential) performance improvement of a pathway-based model is due to the added value of biological information, or instead is a technical artifact. Let a pathway collection (e.g., KEGG) contain *m* pathways *P*_*i*_, *i* = 1, 2, …, *m*, each with *N*_*i*_ genes. Then, a randomly generated pathway collection was produced by randomly assigning *N*_*i*_ genes to pathway *P*_*i*_ . We evaluated the performance of each pathway-based model with multiple randomly generated pathway collections to determine the mean, standard deviation and histogram of the performance metrics of these random pathway baselines.

Finally, the fourth type of baselines correspond to traditional machine learning algorithms, namely random forests (RF). We trained two variations of RF, one with (GEx, T) as input and one with (GEx, FP) as input.

### Cross-dataset analysis by predicting drug responses in CTRPv2 using models trained on GDSC

In addition to the analysis performed using GDSC, we also assessed the generalizability of the deep learning models by performing a cross-dataset analysis. Following the guidelines in a previous study (Sharifi-Noghabi, et al., 2021), we trained the models using area under the dose response curves (AUC) from GDSC dataset to predict AUC of drugs in CTRPv2 (Rees, et al., 2016). For drugs in CTRPv2 dataset, we used their gene expression profile from the cancer cell line encyclopedia (CCLE) (Barretina, et al., 2012). All models were trained using Reactome pathway collection, gene expression and drug targets. Since in this dataset, the gene expressions were quantified using transcript per million (TPM), we also used TPM values for the training set (GDSC). Only common genes between GDSC and CCLE were included. The rest of the preprocessing steps were as described earlier in the manuscript.

## Results

### Models that incorporate KEGG pathway information implicitly outperformed explicit models

Since KEGG was the most commonly used pathway collection in the original studies (Supplementary Table S1), we used the uniform dataset that we formed for this collection to comprehensively evaluate all models. We first focused on GEx data to represent CCLs since all models used GEx modality in their original studies. We also used drug targets to represent compounds since all models could take advantage of this data modality (Morgan fingerprints are not compatible with PathDNN and CDS). Table 3 shows SCC and RMSE values for LCO, LDO, and LPO data splitting strategies (see Supplementary Table S4 for other performance measures and Supplementary Table S5 for statistical tests comparing these models).

**Table 4:** Performance of CDS and PathDSP using KEGG collection, with different input choices. The mean and standard deviations (std) are calculated across cancer cell lines (CCLs). For each method, the input choice that performs best is bold-faced. Since CDS can only use drug targets to represent compounds, the only considered baseline for it is CDS (GEx, T). The following symbols are used in this table: * = original pathway-based model, ▪ = model variant, ↑ = higher value indicates better performance, ↓ = lower value indicates better performance. Supplementary Figure S5 provides visualization of these values in the form of bar plots.

| Model Name | LCO |  | LDO |  | LPO |  |
| --- | --- | --- | --- | --- | --- | --- |
|  | SCC ↑<br>(±std) | RMSE ↓<br>(±std) | SCC ↑<br>(±std) | RMSE ↓<br>(±std) | SCC ↑<br>(±std) | RMSE ↓<br>(±std) |
| PathDSP (GEx, CNV, Mut, FP, T) ★ | **0.883**<br>(±0.044) | 1.308<br>(±0.276) | **0.470**<br>(±0.130) | **2.477**<br>(±0.286) | **0.893**<br>(±0.068) | **1.020**<br>(±0.239) |
| PathDSP (GEx, T) ■ | 0.882<br>(±0.045) | **1.283**<br>(±0.230) | 0.380<br>(±0.146) | 2.648<br>(±0.287) | 0.876<br>(±0.074) | 1.103<br>(±0.256) |
| PathDSP (GEx, FP) ■ | 0.882<br>(±0.043) | 1.297<br>(±0.227) | 0.139<br>(±0.146) | 2.944<br>(±0.327) | 0.887<br>(±0.068) | 1.051<br>(±0.243) |
| CDS (GEx, CNV, T) ★ | 0.777<br>(±0.049) | 1.625<br>(±0.189) | **0.378**<br>(±0.164) | **2.606**<br>(±0.320) | **0.776**<br>(±0.083) | **1.478**<br>(±0.287) |
| CDS (GEx, T) ■ | **0.789**<br>(±0.098) | **1.603**<br>(±0.254) | 0.345<br>(±0.156) | 2.724<br>(±0.317) | 0.769<br>(±0.106) | 1.508<br>(±0.304) |

**Table 5:**
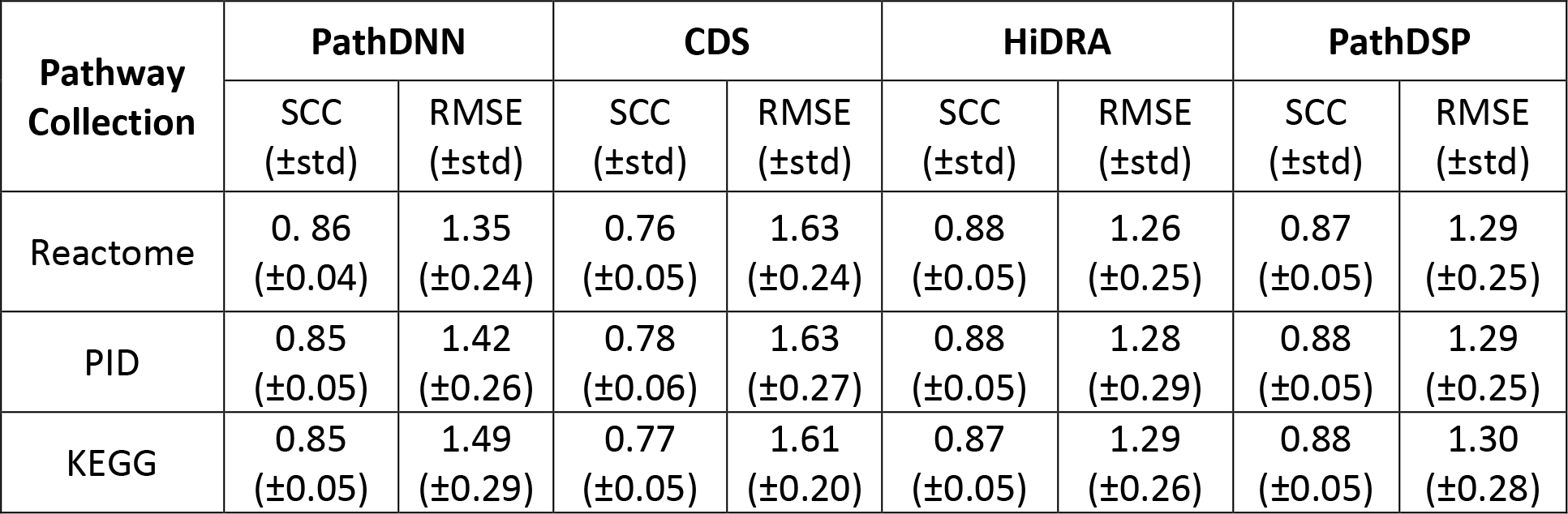
Performance of pathway-based models using different pathway collections. Models with input data used in their original studies are used in this table. More specifically, the models correspond to PathDNN (GEx, T), CDS (GEx, CNV, T), HiDRA (GEx, FP), and PathDSP (GEx, CNV, MuT, FP, T). Mean and standard deviation are calculated across cell lines using the leave-cell- lines-out (LCO) evaluation. Supplementary Figure S12 provides visualization of these values in the form of bar plots and Supplementary Table S9 provides comparison of these models using Wilcoxon signed rank tests.

PathDSP (GEx, T) outperformed all models in LCO and LPO data splitting schemes, while its performance was close to MLP (GEx, T) baseline in the LDO scheme. Compared to the naive predictor, PathDSP (GEx, T) had a better performance in all evaluations, where the highest difference was observed under the LPO validation scheme with 37% lower average RMSE. Overall, the implicit models (HiDRA and PathDSP) outperformed the universal baselines (MLP and naive predictor) for the majority of evaluations, while the explicit models (PathDNN and CDS) did not outperform them in a considerable number of evaluations (Table 3 and Supplementary Table S4).

Next, we assessed the improvement provided by each deep learning method compared to the corresponding naive predictor (Figure 3A). With regards to RMSE, all these models provided improvement for the majority of CCLs in the LPO framework, which is expected since the prediction task in LPO is significantly easier than LCO and LDO. However, in LDO and LCO, many models could not provide a lower RMSE compared to the naive predictor. The improvement was even less in terms of SCC (Figure 3A). However, PathDSP outperformed the naive predictor for the majority of CCLs in all data splitting setups in terms of RMSE and SCC.

**Figure 3:**
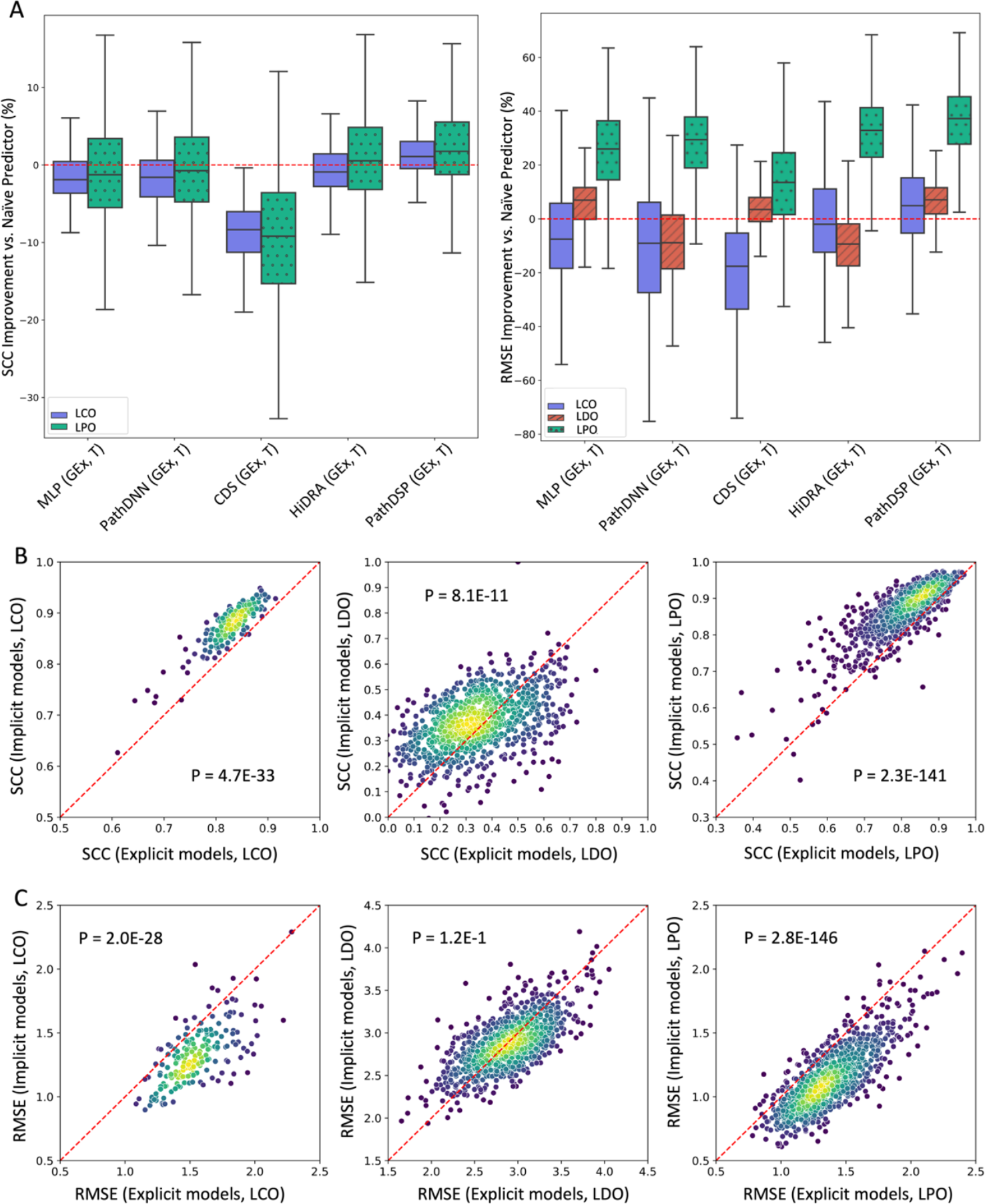
Performance of deep learning models in different data splitting setups. A) The improvement of each model versus naive predictor. Box plots show the distribution of performance improvement for cancer cell lines (CCLs). Each box shows the range between 25^th^ and 75^th^ percentiles, while whiskers show the range of the improvement (excluding outliers). See Supplementary Figure S3 in which the performance improvement of each datapoint (CCL) is also depicted. Spearman’s correlation coefficient (SCC) for naive predictor cannot be calculated in leave-drugs-out (LDO). B-C) Performance of implicit pathway models versus explicit models that use a pathway layer. Each circle represents a CCL. The color of each circle represents the density of circles in its vicinity, where yellow indicates higher density and blue indicates lower density. P-values are calculated using a two-sided Wilcoxon signed-rank test. The average performance of explicit models (PathDNN and CDS) is shown on the x-axis, while the performance of implicit models (PathDSP and HiDRA) is shown on the y-axis. Panel B shows the performance in terms of SCC, while panel C shows it in terms of RMSE. See Supplementary Figure S4 in which the cancer types of CCLs are also depicted.

Next, we sought to directly compare the performance of explicit models against implicit models. For this purpose, we calculated the average performance of the two implicit (PathDSP (GEx, T), HiDRA (GEx T)) and the two explicit (PathDNN (GEx, T), CDS (GEx, T)) models for each CCL, and used a two-sided Wilcoxon signed rank test to assess if one strategy outperforms the other (Figure 3B). Based on SCC, the implicit strategy significantly outperformed the explicit strategy that utilizes a pathway layer, for all three data splitting strategies. A similar pattern was observed using RMSE, but for LDO strategy the difference was not statistically significant. These results further confirm the observation that utilizing an explicit pathway layer does not seem to perform well in prediction of drug response. Supplementary Figure S4 also shows similar scatter plots in which the cancer types of cell lines are depicted, which does not suggest a cancer type-specific pattern.

### Morgan fingerprints of compounds were more informative than drug targets for predicting response of unseen cell lines

Since three of the considered deep learning models (MLP, HiDRA, and PathDSP) can utilize both drug targets (T) and Morgan fingerprints (FP) to represent drugs, we sought to determine which compound representation is most informative for drug response prediction. As can be seen in Figure 4, in all three models, using FP to represent compounds in most cases was superior in terms of SCC in predicting unseen CCLs (LCO) or in predicting unseen CCL-drug pairs (LPO) (Two-sided Wilcoxon signed-rank P<0.05, except for PathDSP LCO). On the other hand, in all three models drug targets were more informative in predicting the response of unseen drugs (LDO). However, one should note that none of the three models performed very well in the LDO data splitting setup and more informative compound representations (e.g., transcriptomic changes in response to compounds (El Khili, et al., 2022) or DL models that directly learn compound representations (Zagidullin, et al., 2021)) may be necessary for such an application to allow generalization to new compounds.

**Figure 4:**
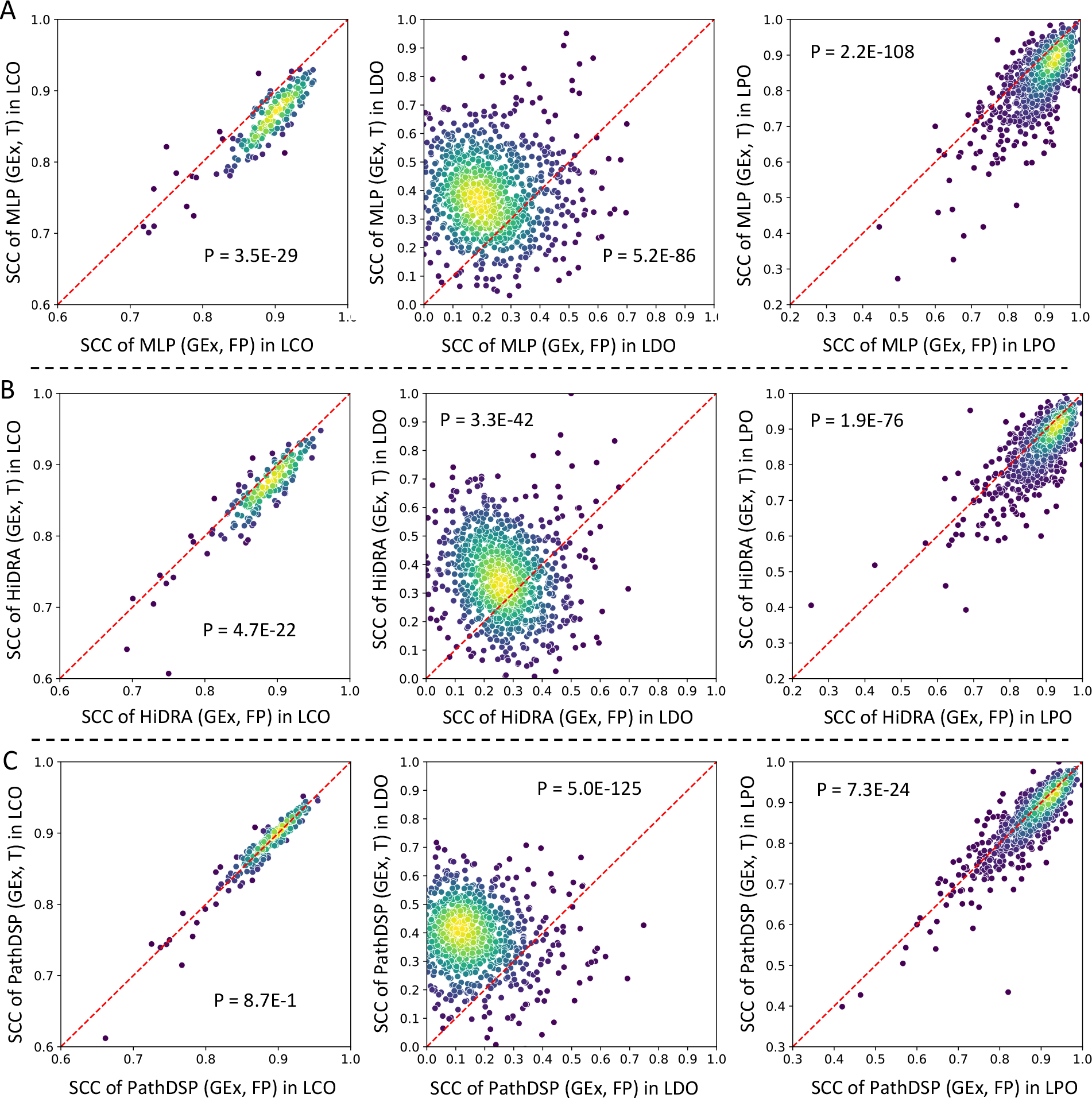
Performance of three models when using drug targets or Morgan fingerprints (FP) to represent drugs in terms of Spearman’s correlation coefficient (SCC). Each circle represents a cancer cell line (CCL). The color of each circle represents the density of circles in its vicinity, where yellow indicates higher density and blue indicates lower density. P-values are calculated using a two-sided Wilcoxon signed-rank test. For all models, the mean SCC when using FP was higher in leave-pairs-out (LPO) and leave-cell-lines-out (LCO), and lower in leave-drugs-out (LDO) compared to when using drug targets (T). A) Performance of MLP (GEx, T) versus MLP (GEx, FP). B) Performance of HiDRA (GEx, T) versus HiDRA (GEx, FP). C) Performance of PathDSP (GEx, T) versus PathDSP (GEx, FP). Only models that could utilize both FP and T to represent drugs were used for this analysis.

It is worth noting that although PathDSP (GEx, T) and HiDRA (GEx, T) both outperformed the MLP (GEx, T) baseline in LCO and LPO evaluation, MLP (GEx, FP) baseline outperformed all other models, independent of which CCL or drug representations they used in both LCO and LPO (Supplementary Table S4). This is an important observation that shows that a simple MLP baseline, when used with appropriate inputs could achieve comparable or better results compared to various interpretable models. This observation is concordant with the observation in (Kuenzi, et al., 2020), where the authors found that the interpretable version of their model resulted in comparable performance to the matched black-box model. Interestingly, RF (GEx, FP) provided the best performance in terms of SCC and RMSE in LCO, showing that sometimes traditional machine learning methods can achieve similar or better results compared to deep learning methods, an observation also made in (Chen and Zhang, 2022).

### Integrating multiple data modalities improves performance of PathDSP and CDS

Among pathway-based methods that we considered in this study, two of them (CDS and PathDSP) used multiple data modalities in their original study (Supplementary Table S1). Table 4 compares the performance of these methods when data modalities chosen by the original study were used as inputs against their performance when only GEx was used (see Supplementary Table S6 for comparison of these models using two-sided Wilcoxon signed rank tests). The original PathDSP model uses GEx, somatic mutation, and CNV as CCL features, as well as Morgan fingerprints and drug targets as compound features, which for clarity we denote as PathDSP (GEx, CNV, MuT, FP, T). PathDSP (GEx, CNV, MuT, FP, T) outperformed both PathDSP (GEx, T) and PathDSP (GEx, FP) in 5 out of 6 evaluations (all except RMSE in LCO, Table 4). The original CDS model uses GEx and CNV as CCL features and drug targets as compound features, which for clarity we denote as CDS (GEx, CNV, T). The original CDS (GEx, CNV, T) model also outperforms CDS (GEx, T) in all evaluations except for LCO approach. Overall, these results suggest that using multiple data modalities can improve the performance of each model. However, it is important to remind that MLP (GEx, FP) outperformed all models (including the multi-modality versions of PathDSP and CDS) in LCO and LPO evaluations (Supplementary Table S4).

### Randomly generated pathways provide comparable results to biological pathway collections for prediction of drug response in unseen cell lines

Next, we sought to determine whether the performance of pathway-based models can be attributed to the biological information in the pathways, or if randomly generated pathways can also result in a similar performance. For this purpose, we randomly assigned genes to pseudo- pathways while matching the size of the pathways in the KEGG collection. Figure 5A shows the percentage of improvement in the form of a heat map, where we compared the original pathway- based models (PathDNN, CDS, HiDRA, PathDSP) and their model variants with their corresponding random pathway baselines. Figure 5B and Supplementary Figures S6-S10 show the distribution of SCC and RMSE of different models for each randomly generated pathway collection in different validation schemes.

**Figure 5:**
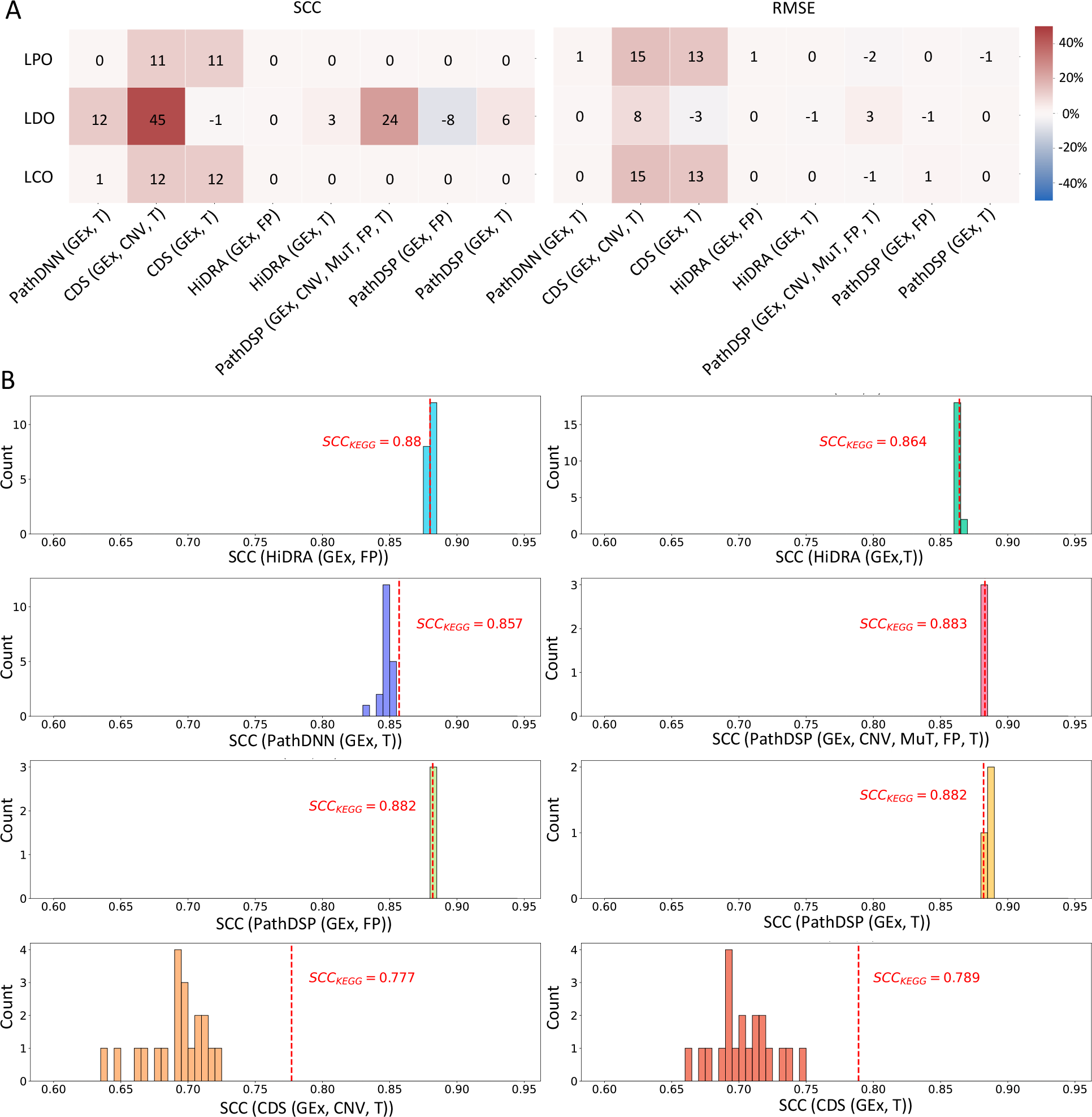
Performance of pathway-based models using KEGG or randomly generated pathways. 1) Percentage of improvement (or deterioration) of different models when using KEGG compared to their mean performance when using randomly generated pathways. B) The histograms show the distribution of mean Spearman’s correlation coefficient (SCC) of random pathway baselines using the leave-cell-lines-out (LCO) validation scheme. Vertical dashed red lines show SCC of the model when using KEGG pathways. Twenty random pathway baselines were constructed for each model, except for PathDSP models. Since PathDSP requires 1000 permutation tests for each type of input data, only three random pathway baselines were constructed due to its extremely high computational requirement.

As can be seen in Figure 5, in the LCO and LPO evaluations, biological pathways provided almost no improvement compared to their randomly generated counterparts (percentage of improvement between -2% and 2%) for PathDSP, PathDNN, and HiDRA. CDS was the only exception for which an improvement of up to 15% was obtained using biological pathways (specifically for the original CDS (GEx, CNV, T) model). While it is difficult to conclusively determine why CDS benefits from biological pathways, our conjecture is that this is due to its unique architecture combined with the use of CNV input data. Moreover, it is important to note that despite this improvement, CDS (GEx, CNV, T) and CDS (GEx, T) had much worse performance compared to the other models (Figure 5B and Table 3).

In the LDO evaluation, the two models that used Morgan fingerprints to represent compounds, PathDSP (GEx, FP) and HiDRA (GEx, FP) did not perform better with biological pathways compared to randomly generated pathways. On the other hand, the majority of models that used drug targets experienced an improvement compared to randomly generated pathways. We investigated this behavior further by inspecting the number of drug targets in KEGG pathways and the randomly generated pathways (Supplementary Figure S11). Comparing the number of targets in each KEGG pathway with the randomly generated pathways of the same size showed that in the majority of pathways (230 out of 332 pathways, approximately 70%), the number of drug targets in the KEGG pathways were larger. Since drug targets are integrated with pathway information to obtain drug embeddings, this difference in the number of drug targets results in less informative and less distinguishable drug embeddings in the case of randomly generated pathways. For example, in PathDNN where drug targets are represented as binary features, the random pathway nodes are connected to many zero-valued drug features. Such nodes do not participate much in capturing the similarities or differences of drugs, leading to embeddings that are not as informative as their biological pathway counterparts in capturing patterns of similarity and dissimilarity of drugs. This observation is also in line with a recent study that showed better predictions could be obtained for compounds with diverse target classes (Kuenzi, et al., 2020). The issue mentioned above is particularly important in the case of LDO, since unlike LCO and LPO where all drugs in the test set have been seen by the model during training, the model must learn drug similarity/dissimilarity patterns in order to make predictions for new drugs not observed during training. This results in a deterioration of performance in random-pathway models (parituclarly in LDO) compared to their biological counterparts observed in Figure 5.

### Effect of pathway collection choice on drug response prediction

We next sought to investigate which pathway collection is more suitable for the drug response prediction task. For this purpose, we compared the performance of each pathway-based model (in their original architecture and using original input features) using each of these collections (see Supplementary Tables S7 and S8 for the performance of all models and their variants using PID and Reactome). To ensure a fair comparison, we only included (drug, CCL) pairs in the test sets that were shared among all three uniform datasets. We used the LCO data splitting approach, since the overlap among the test samples of the three uniform datasets was largest in this strategy (21525 pairs versus 3277 in LDO and 851 in LPO).

Table 5 and Supplementary Figure S12 show the mean and standard deviation of SCC and RMSE of each model using all three pathway collections. Overall, we observed that the performance of most models did not vary drastically based on the choice of pathway collection. However, Reactome pathway provided slightly better results for the majority of the methods, being the top performing option for 3 (out of 4 methods) based on RMSE. We hypothesized that this is due to the larger number of pathway annotations included in this database for our use-case (1608 pathways in Reactome compared to 332 in KEGG and 196 in PID), resulting in a more comprehensive representation of the input data.

To test whether the large number of pathways in Reactome can explain its better performance, we randomly downsampled the pathways in this collection. Figure 6 shows the SCC for PathDNN (GEx, T) using different number of pathways removed (x-axis). We focused on PathDNN (GEx, T), since it achieved its best performance when using Reactome pathway collection (compared to PID or KEGG). For each value on the x-axis, downsampling was performed ten times and the results were used to calculate the mean and standard deviation in the LCO setup (Supplementary

**Figure 6:**
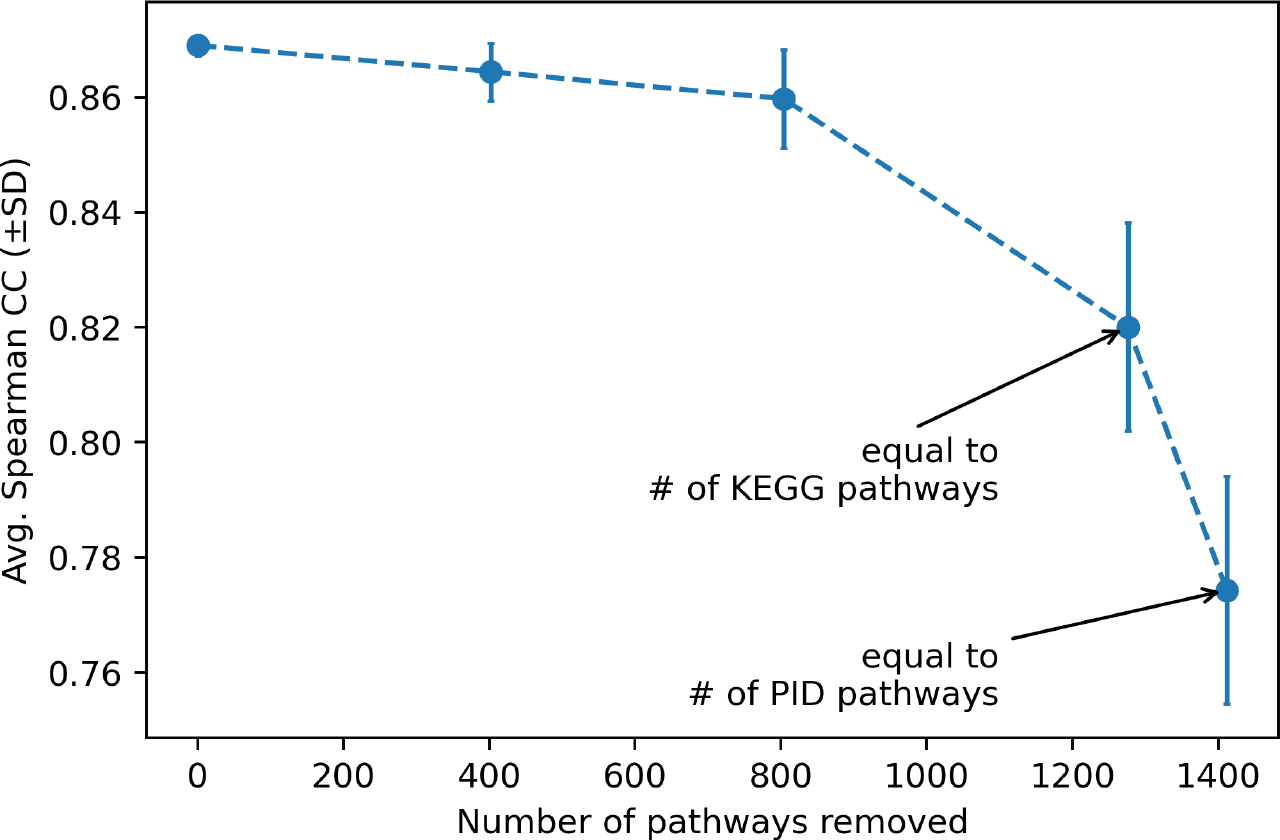
Performance of PathDNN (GEx, T) with downsampled Reactome pathways. The y-axis shows the mean (Avg.) and standard deviation (SD) of Spearman’s correlation coefficient (SCC) and the x-axis shows the number of pathways removed from the Reactome collection.

Table S10 provides details of each run). This figure shows that indeed, the number of pathways in the Reactome collection plays a major role in its performance: as more pathways are removed, the performance of PathDNN (GEx, T) deteriorates, with the lowest mean SCC value obtained when only 196 (equal to the number of pathways in PID) have remained. This signifies that the comprehensiveness of Reactome has enabled PathDNN to achieve better results. Interestingly, the performance of this model with PID or KEGG was much better compared to the downsampled version of Reactome with the same number of pathways (Table 5 and Figure 6). We attribute this to the increasing probability of removing an important pathway during random downsampling of Reactome, as well as the quality of the curated pathways in KEGG and PID.

### Cross-dataset performance of pathway-based models

In addition to the analysis performed using GDSC reported earlier, we also assessed the generalizability of the deep learning models to predict response of drugs in CTRPv2 (Rees, et al., 2016). For this purpose, we trained the models on GDSC using drug AUC values and assessed their performance on the prediction of AUC of drugs in CTRPv2. For consistency, all models were trained using gene expression and drug targets. Supplementary Table S11 shows the results in various data splitting and evaluation setups. Similar to our previous analyses on GDSC, in LCO and LPO setup, implicit models performed better compared to explicit models and also outperformed MLP. However, in the LDO setup MLP baseline achieved the best performance. The performance of all models on CTRPv2 deteriorated compared to their performance on GDSC, highlighting the challenging nature of this task.

## Discussion

Recently, several deep learning methods have been proposed to enable a higher interpretability of drug response prediction and to improve the prediction performance. In this study, we set out to investigate four methods that try to achieve these goals by incorporating pathway information under various validation schemes and answer five important questions discussed earlier. The models were tested to predict drug response for unseen (CCL, drug) pairs, unseen CCLs, and unseen drugs. We compared these methods against four types of baseline models, two of which were usually overlooked in previous studies.

First, we observed that models that incorporate a dedicated explicit pathway layer and connect gene nodes in a previous layer to pathways based on pathway membership perform worse compared to models that implicitly (e.g., using attention mechanisms or pathway enrichment scores) incorporate pathway information. In fact, in many occasions explicit models’ performance was inferior to a black-box simple MLP model with similar input. This suggests that direct encoding of gene-pathway membership is not an effective strategy to incorporate pathway information. The overly sparse connections between the gene and pathway layer may be the cause for the unsatisfactory performance of these methods (due to a reduction in their capacity), supported by the observation that their MLP counterparts (with fully connected layers) achieved a better performance. Another limitation of explicit models is that they can only utilize gene-level drug representations, limiting usable drug features to drug targets. Our analysis using methods that could utilize both drug targets and Morgan fingerprints showed the latter to be superior in prediction of response for unseen CCLs or unseen pairs. However, recent studies have suggested that alternative drug representations such as transcriptomic changes in response to compounds (El Khili, et al., 2022) or DL-based fingerprints (Zagidullin, et al., 2021) may improve performance of drug response predictors.

Our analyses also showed that while implicit models generally performed better in predicting unseen CCLs and unseen pairs, a comparable performance can be achieved when instead of biological pathways, randomly generated pathways are used. Moreover, in these validation setups a black-box MLP that used Morgan fingerprints for drug representation outperformed all pathway-based models. Put together, these results suggest that to make the models interpretable, these approaches inevitably make assumptions that cannot fully capture the nuances of drugs’ mechanisms of action in cancer cell lines, resulting in comparable or worse performance compared to black-box models.

Our analyses also allowed us to assess the difficulty of drug response prediction in different setups. While at first glance, Table 3 may suggest that predicting response of unseen drugs are much more challenging than unseen CCLs, a more appropriate comparison can be made using Figure 3A, where the models’ performance improvement under each validation setup was compared against a naive predictor. This figure shows that predicting response of unseen pairs is much easier compared to prediction for unseen CCLs and unseen drugs. This is not surprising, since this is a transductive setup and drugs and CCLs in the test set are present in the training set (but not together). This setup is useful for imputation of missing drug response values but cannot be used to predict response to new CCLs or new drugs. On the other hand, predicting response of unseen drugs and unseen CCLs are much more difficult and most models cannot provide a better prediction for the majority of CCLs in these two setups (Figure 3A). Moreover, the performance of all models deteriorated when used to predict the drug response in a different dataset (CTRPv2), revealing the challenging nature of this task.

The contrast between conclusions one may draw from Table 3 and Figure 3A (discussed above) demonstrates the potential for obtaining inflated performance measures in the LCO framework, in which for a given CCL in the test set, the log IC50 value of different drugs are to be predicted. Supplementary Figure S13 shows the distributions of log IC50 values for each drug (across CCLs, panel A) and each CCL (across drugs, panel B) in our dataset. While these values vary across both drugs and CCLs, the identity of a drug plays a bigger role in determining its log IC50 value compared to the identity of the CCL to which it was administered (i.e., log IC50 values are more drug-specific than CCL-specific). Supplementary Figure S13C better clarifies this point by depicting the histogram of false discovery rates (FDRs) obtained from comparing the local distribution of log IC50 values per drug (purple) or per CCL (green) and the global distribution of the log IC50 values using Mann-Whitney U tests. Although for the majority of drugs (93%) drug-specific log IC50 values (across all CCLs, but for a specific drug) are significantly different (FDR < 0.05) from the global log IC50 values (across all drugs and CCLs), that is true for only 43% of CCLs. This implies that by simply knowing the identity of a drug, a model can rank different drugs based on their log IC50 values for an unseen CCL rather well. To overcome this issue and avoid reporting unrealistically inflated metrics, one should focus on improvement compared to a naive predictor (an approach that we adopted in this study), or should normalize log IC50 values of each drug across CCLs to make them comparable to each other (an approach that we adopted in (Hostallero, et al., 2022)).

We also compared the performance of different models using three pathway collections, PID, KEGG, and Reactome. Even though there was not a major difference in the performance of models when substituting one collection for the other, Reactome collection resulted in slightly better performance. This seems to be due to the larger number of pathways in this collection compared to the other two. However, since randomly generated pathway collections also provided comparable performance based on the models considered in this study, it is not possible to draw a conclusive determination regarding which pathway collection may be more useful for the drug response prediction task.

This study focused on evaluating the effect of incorporating pathway information from the perspective of model performance and we did not evaluate these models based on their level of interpretability. A study that focuses on interpretability aspect of these models would be very insightful and complementary to the current study. For example, one can take a closer look at the feature attributions of these pathway-based models using explainers such as DeepLIFT (Deep Learning Important FeaTures) (Shrikumar, et al., 2017), CXPlain (Schwab and Karlen, 2019), and SHAP (Shapley Additive exPlanations) (Lundberg and Lee, 2017) to estimate feature importance and identify genes or other biological features that have substantial influence on the model predictions. Such analysis can be done for all pathway-based models to check if the most important/predictive sub-networks or the top contributing genes extracted from each model have any overlap. If such overlap exists between the pathway-based models, further studies can be done by validating the findings with existing literature or conducting experiments under a lab setting. Analysis on model interpretability will complement the insights obtained from model performance evaluation and together provide a more holistic view for the effect of pathway incorporation on drug response prediction.

In conclusion, we believe that while interpretability is a very crucial aim in precision medicine, new models are necessary to enable a higher degree of interpretability while at the same time improve the drug response prediction performance. In addition, it is not sufficient for these models to show a better performance compared to their black-box counterparts, and they need to also evaluate their models against randomly generated pathways (with similar pathway sizes to the original collection) and naive predictors to control for different types of biases.

### Data and Code Availability

Input data for the evaluated models is provided at https://zenodo.org/record/7101665#.YzS79HbMKUk. The implementation of the models are available at https://github.com/Emad-COMBINE-lab/InterpretableAI_for_DRP.

## Supporting information

Supplementary File S1 containg Supplementary Figures

