## Supplementary File S1 containg Supplementary Figures for "Interpretable deep learning architectures for improving drug response prediction performance: myth or reality?"

**Supplementary Methods, Figures and Tables for "Interpretable deep learning architectures for improving drug response prediction performance: myth or reality?"**

Yihui Li[1], David Earl Hostallero[1,2], and Amin Emad[1,2,3,*]

[1] Department of Electrical and Computer Engineering, McGill University, Montreal, QC, Canada
[2] Mila, Quebec AI Institute, Montreal, QC, Canada
[3] The Rosalind and Morris Goodman Cancer Institute, Montreal, QC, Canada

* Corresponding Author:
Amin Emad
755 McConnell Engineering Building
3480 University Street, Montreal, Quebec, Canada, H3A 0E9


**Supplementary Methods:**

**Model Description**

Here, we provide an overview of different models considered in the study. For a detailed description of the architectures, hyperparameter tuning (including the selected values), and training see Supplementary File S2.

**PathDNN (explicit pathway integration model)**

PathDNN is a pathway-guided deep neural network that consists of four hidden layers, of which the first hidden layer is dedicated to representing pathways. All other layers are fully-connected, except for the input and pathway layers. The connections between the input and pathway layers are defined based on pathway membership, therefore only member genes of each pathway are considered during the model learning process.

We evaluated the original PathDNN model and its random pathway baseline (denoted by PathDNN rand). PathDNN rand has the same architecture and pipeline as the original model, except the gene-pathway connections (determined by the mask matrix $M$) are defined randomly. To generate these random connections, we fixed the total number of member genes $N_i$ for each pathway $P_i$ and selected $N_i$ number of random genes. We repeated this process twenty times to generate twenty random gene-pathway mask matrices, thus obtaining twenty different random pathway baselines. We then calculated the mean performance metrics of these twenty baselines to benchmark the original PathDNN model that uses biological pathway information.

**ConsDeepSignaling (CDS) (explicit pathway integration model)**

ConsDeepSignaling (CDS) is another deep learning model constrained by pathways. It contains six hidden layers, where the first two hidden layers are used to represent genes and pathways. The input layer consists of nodes representing gene expression (GEx), copy number variation (CNV), and drug target (T) information. The trio of input feature nodes are connected to their corresponding gene node in the proceeding gene layer. The connections between the gene and pathway layer are determined by pathway membership, which is the same approach used by PathDNN.

Because the original CDS model uses multi-omics data, we tested a single-omic variant of CDS (named CDS (GEx, T), where short names in the bracket represent the types of cell line and drug features used by the model) to ensure fair comparison with other single-omic models evaluated in this study. This model variant only uses gene expression data as cell line features, instead of both gene expression and CNV data like the original model. For both CDS and CDS (GEx, T), corresponding random pathway models are constructed in a similar fashion as PathDNN rand.

**HiDRA (implicit pathway integration model)**

HiDRA is a hierarchical network for drug response prediction with incorporated attention mechanism. Unlike PathDNN and CDS, its operation does not involve the use of a pathway layer. The pathway information is integrated into the model using gene-level attention module for the member genes of each pathway, from which pathway activation scores are determined. These activation scores are then taken by a pathway-level attention module to generate weighted pathway importance scores. Finally, drug features are concatenated with the learned pathway importance scores and fed into a two-layer MLP to output the final drug response prediction.

The original HiDRA model uses gene expression data as cell line features and Morgan fingerprints (FP) as drug features. In addition to evaluating the original model, we also tested a model variant named HiDRA (GEx, T), which uses target data as drug features. The random pathway models for HiDRA and HiDRA (GEx, T) are constructed using randomly selected genes for the gene-level attention module. We constructed twenty random pathway baselines for each of HiDRA and HiDRA (GEx, T), then the mean performance metrics are determined to benchmark the performance of models that use actual pathway information.

**PathDSP (implicit pathway integration model)**

The architecture of PathDSP is like that of an MLP. The novelty of this model comes from the strategy used to transform molecular features into pathway-level features. The original study performed pathway enrichment analysis on gene expression, CNV, somatic mutation, and drug target data. The final pathway-level features are matrices of pathway enrichment scores specific to every pathway and with respect to each cell line or drug. Despite the different data types, all molecular features are transformed onto the same level of representation using pathway enrichment analysis. These pathway enrichment scores of the different types of features are then concatenated to form the input to the model. As a result, the pathway information is implicitly embedded in the input data to the model.

The original PathDSP uses five different types of input features (gene expression, somatic mutation, CNV, drug structural data (Morgan fingerprints), drug target data). In order to compare PathDSP fairly with other models, we evaluated two single-omic variants of PathDSP, namely PathDSP (GEx, FP) and PathDSP (GEx, T), which use gene expression as cell line features and Morgan fingerprints, drug target data as drug features, respectively. The pathway enrichment analysis that generates pathway enrichment scores for performs 1000 permutation tests for each type of input data and therefore is very computationally expensive. Due to this computational limitation, we were only able to construct three (as opposed to twenty for the other pathway-based models) random pathway baselines for PathDSP and its model variants.

### Supplementary Tables

**Supplementary Table S1:** Summary of input data and pathway databases used by original studies of pathway-based models. CCL = cancer cell line, GEx = gene expression, Mut = somatic mutation, CNV = copy number variation, T = drug target data, FP = Morgan fingerprint (drug structural data).

| Model Name | CCL | Drug | Num. CCLs | Num. Drugs | Pathway |
|---|---|---|---|---|---|
| PathDNN [1] | GEx | T | 970 | 250 | KEGG [2] |
| CDS [3] | GEx, CNV | T | 791 | 24 | KEGG [2] |
| HiDRA [4] | GEx | FP | 968 | 235 | KEGG [2] |
| PathDSP [5] | GEx, CNV, Mut | T, FP | 319 | 153 | PID [6] |

**Supplementary Table S2:** Data modality and source of the uniform datasets.

| Data Modality | Source |
|---|---|
| Gene Expression (GEx) | GDSC [7] |
| Somatic Mutation (Mut) | GDSC [7] |
| Copy Number Variation (CNV) | GDSC [7] |
| Drug Structural Data (FP) | PubChem [8] |
| Drug Target Data (T) | STITCH [9] |
| Protein-Protein-Interaction (PPI) Network | STRING Experimental PPI [10] |
| Pathway Data | KEGG [2], PID [6], Reactome [11] |

**Supplementary Table S3:** Information on the identity and characteristics of cell lines and drugs used in this study for the uniform dataset corresponding to each pathway collection. The results are provided as a separate xlsx file.

**Supplementary Table S4:** Summary of results for the uniform dataset generated for KEGG pathway collection. The results are provided as a separate xlsx file.

**Supplementary Table S5:** Pairwise comparison of methods shown in Table 3 using a two-sided Wilcoxon signed rank test. The results are provided as a separate xlsx file.

**Supplementary Table S6:** Pairwise comparison of methods shown in Table 4 using a two-sided Wilcoxon signed rank test. The results are provided as a separate xlsx file.

**Supplementary Table S7:** Summary of results for the uniform dataset generated for PID pathway collection. The results are provided as a separate xlsx file.

**Supplementary Table S8:** Summary of results for the uniform dataset generated for Reactome pathway collection. The results are provided as a separate xlsx file.

**Supplementary Table S9:** Pairwise comparison of methods shown in Table 5 using a two-sided Wilcoxon signed rank test. The results are provided as a separate xlsx file.

**Supplementary Table S10:** The effect of downsampling on the performance of PathDNN (GEx, T) using Reactome pathway collection in the leave cell lines out setup. The results are provided as a separate xlsx file.

**Supplementary Table S11:** The cross-dataset generalizability performance of models on predicting drug response in the CTRPv2 database using models trained on the GDSC database. Reactome pathway collection was used for all models. The results are provided as a separate xlsx file.

**Supplementary Figures:**

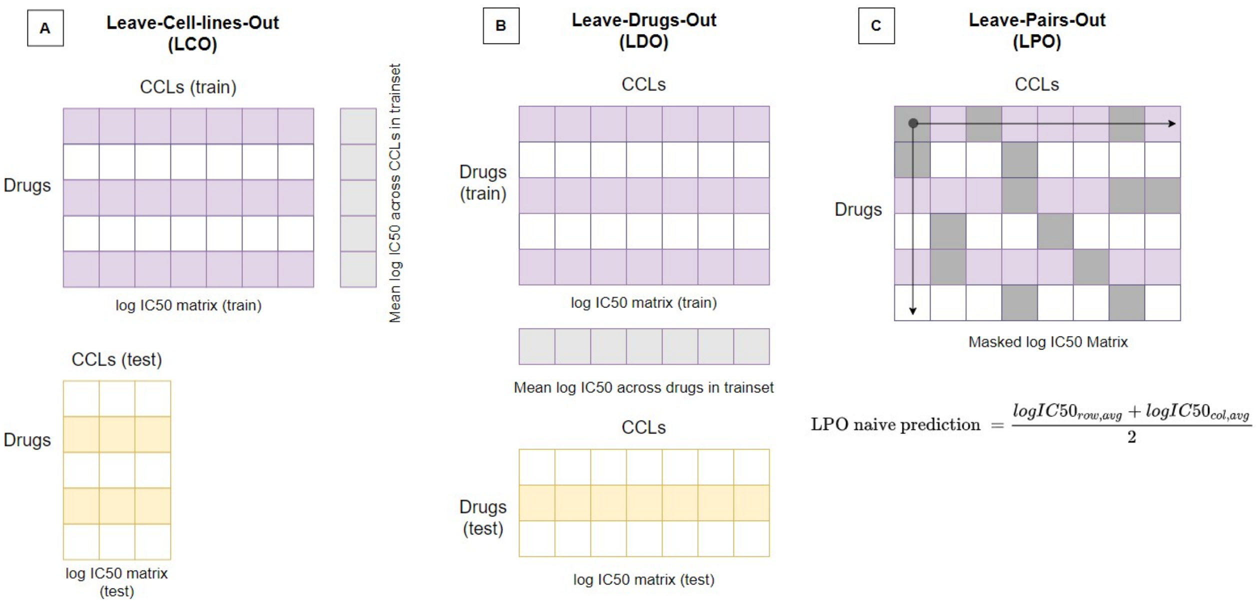


**Figure S1:** The drug response data is first divided into training and test set using different validation schemes (LCO, LDO, LPO). The divided data folds are then used by the naive predictor to generate predictions. A) In the LCO setup, the naive predictions are generated by calculating average log IC50 values of each drug across CCLs in the training set (shown by purple matrix). The obtained naive predictions are drug-specific and independent of CCLs. This vector of naive predictions can then be compared with the ground-truth log IC50 values in the test set (shown by yellow matrix) to obtain performance metrics. B) In the LDO setup, naive predictions are obtained in a similar fashion as the LCO setup, except that the average log IC50 values are calculated across all drugs in the training set. C) In the LPO setup, the log IC50 matrix is first masked to only keep samples in the training set (masked elements are shown in gray). Then for each sample in the test set, average log IC50 values are calculated by row (across CCLs) and column (across drugs). The LPO naive prediction is reported as the average of the row-wise and column-wise mean log IC50 values.

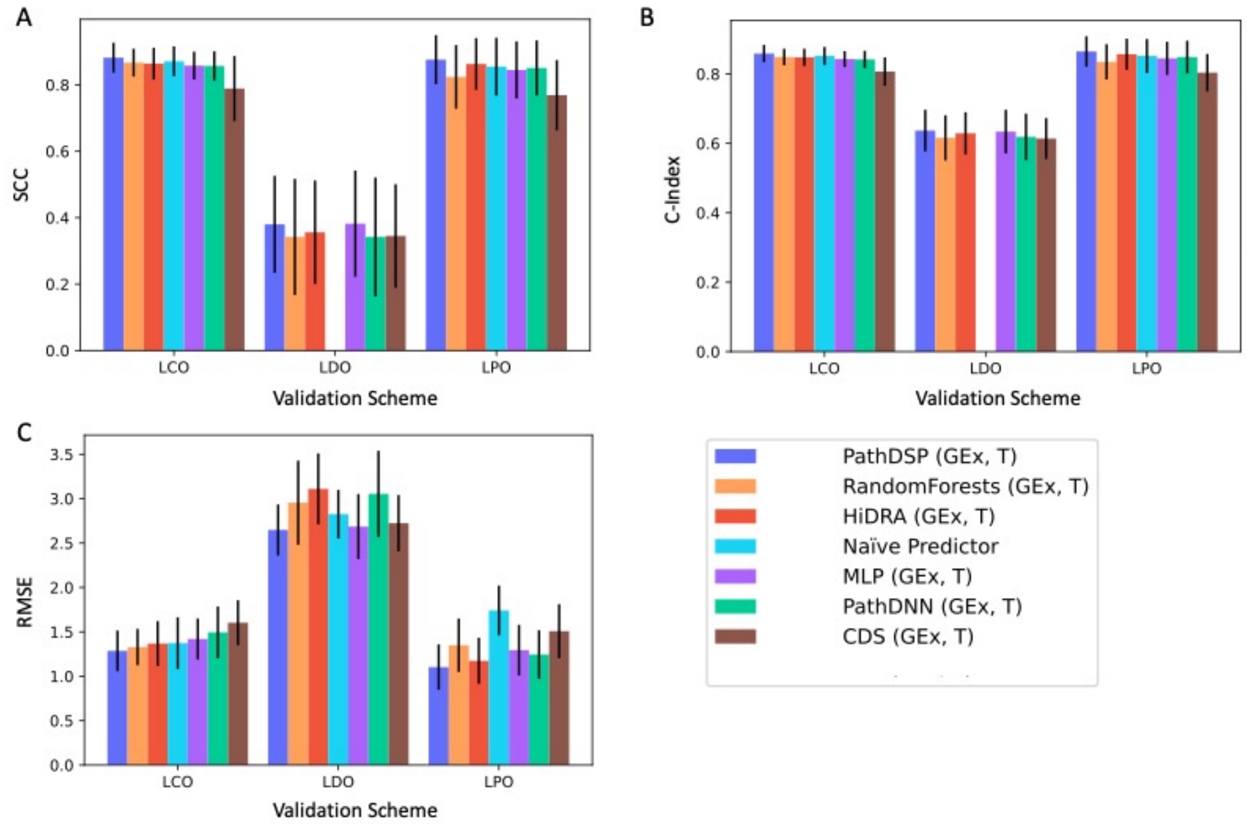


**Figure S2:** Performance of different models using KEGG pathway collection, with gene expression (GEx) and drug targets (T) as inputs. The mean and standard deviations are calculated across CCLs. Standard deviations are shown as error bars. A) Spearman Correlation Coefficient (SCC), B) Concordance Index (C-Index), C) Root Mean Squared Error (RMSE). The LDO SCC and C-Index cannot be calculated for the naive predictor since in this case it outputs the same value for all CCLs. This figure corresponds to Table 3 in the main manuscript.

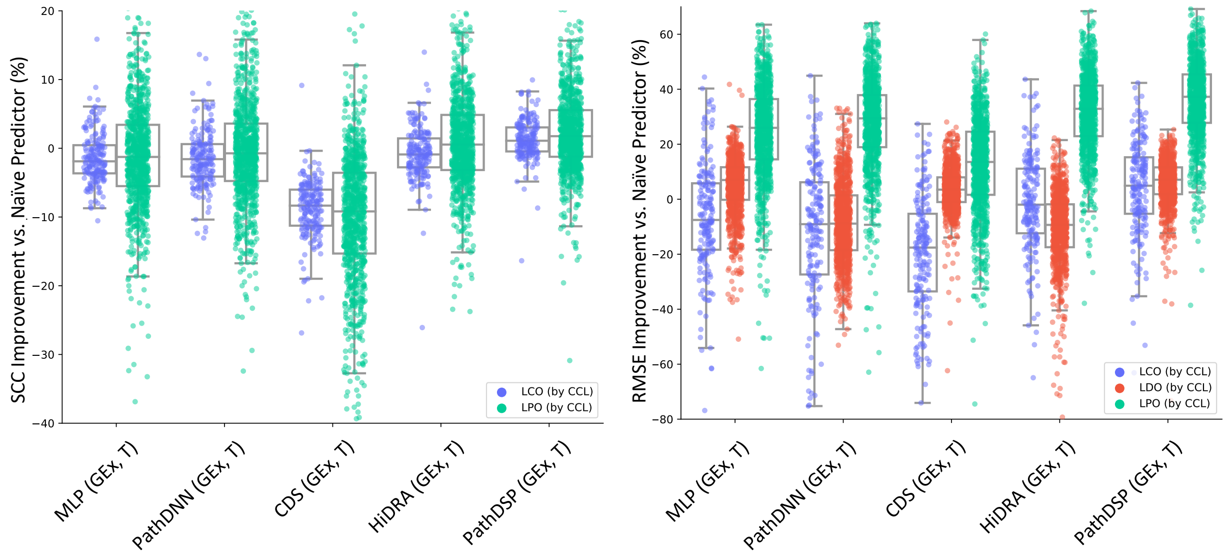


**Figure S3:** The improvement of each model versus naïve predictor. Box plots show the distribution of performance improvement for CCLs (depicted as circles). Each box shows the range between 25$^{th}$ and 75$^{th}$ percentiles, while whiskers show the range of the improvement (excluding outliers). SCC for naïve predictor cannot be calculated in LDO.

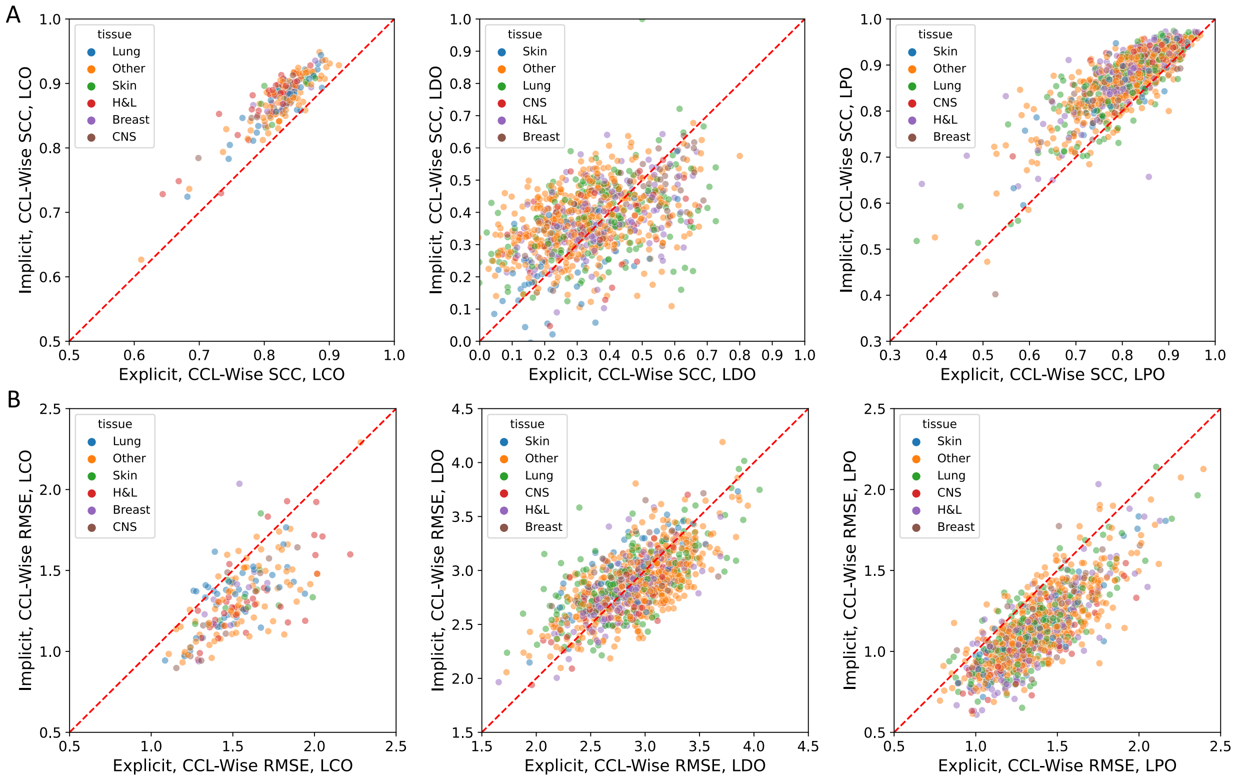


**Figure S4:** Performance of implicit pathway models versus explicit models that use a pathway layer based on CCL tissue of origin. Each circle represents a CCL. The average performance of explicit models (PathDNN and CDS) is shown on the x-axis, while the performance of implicit models (PathDSP and HiDRA) is shown on the y-axis. Panel A shows the performance in terms of SCC, while panel B shows it in terms of RMSE. Top 5 tissues are selected based on number of CCLs in the test set (CNS = Central Nervous System, H&L = Haematopoietic and Lymphoid, Other = all other tissue types).

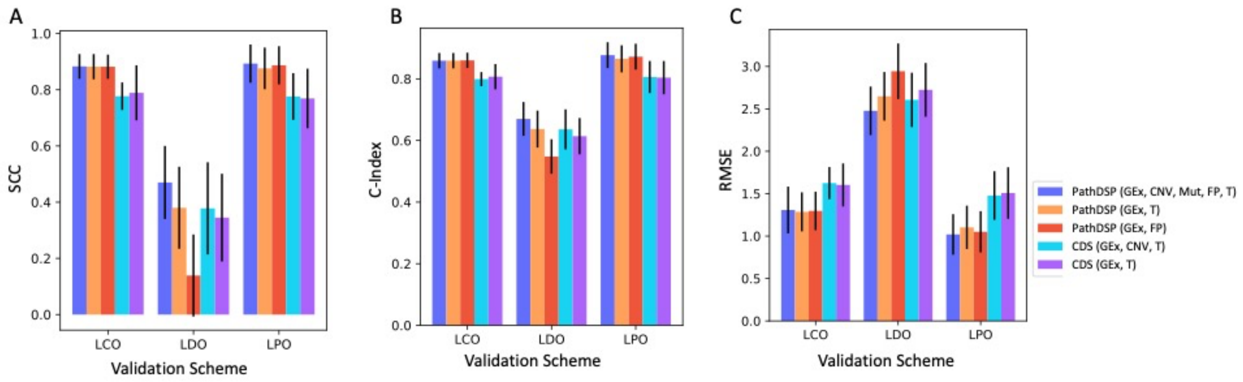


**Figure S5:** Performance of CDS and PathDSP using KEGG collection, with different input choices. The mean and standard deviations are calculated across CCLs. Standard deviations are shown as error bars. A) Spearman Correlation Coefficient (SCC), B) Concordance Index (C-Index), C) Root Mean Squared Error (RMSE). This figure corresponds to Table 4 in the main manuscript.

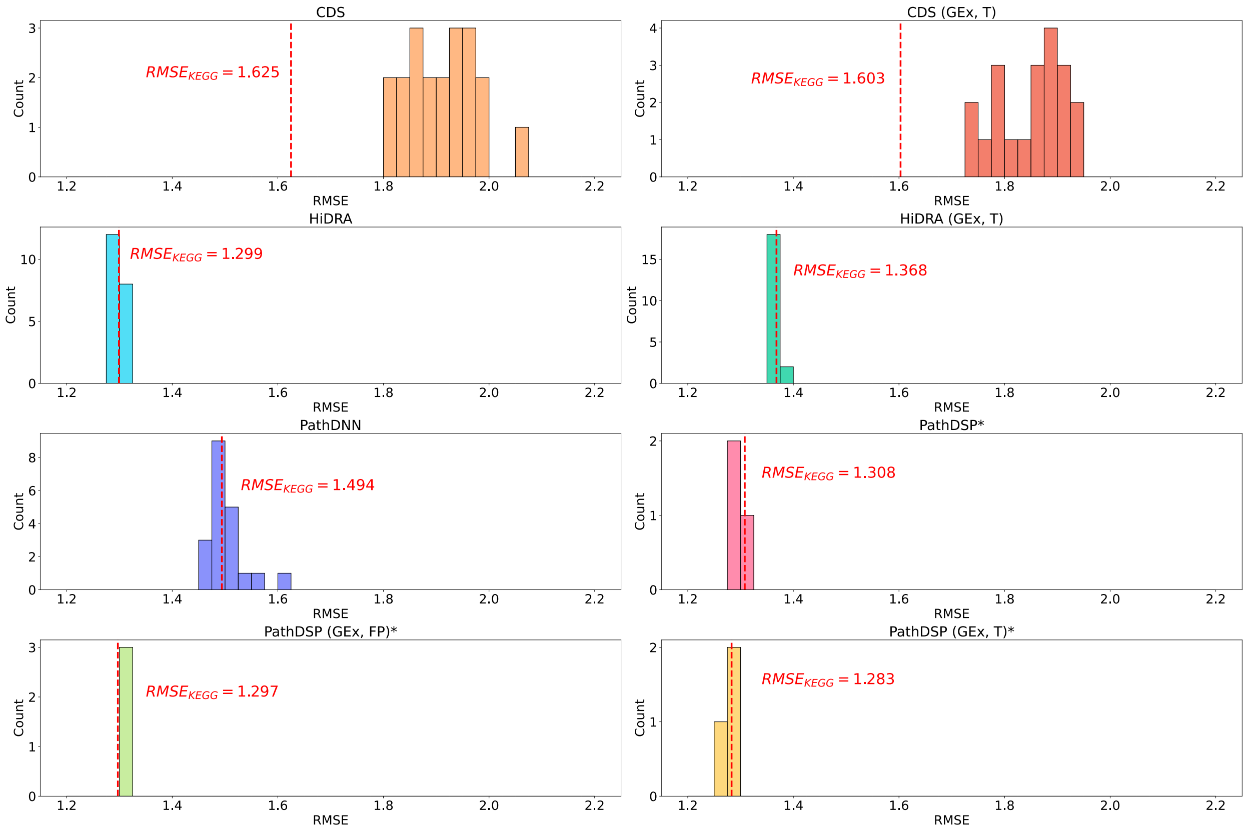


**Figure S6:** Performance of pathway-based models using KEGG or randomly generated pathways. The histograms show the distribution of mean RMSE of random pathway baselines using the LCO validation scheme. Vertical dashed red lines show RMSE of the model when using KEGG pathways. In this figure, CDS corresponds to CDS (GEx, CNV, T), HiDRA corresponds to HiDRA (GEx, FP), PathDNN corresponds to PathDNN (GEx, T), and PathDSP corresponds to PathDSP (GEx, CNV, MuT, FP, T) in Table 2. *Since PathDSP requires 1000 permutation tests for each type of input data, only three random pathway baselines were constructed due to its extremely high computational requirement.

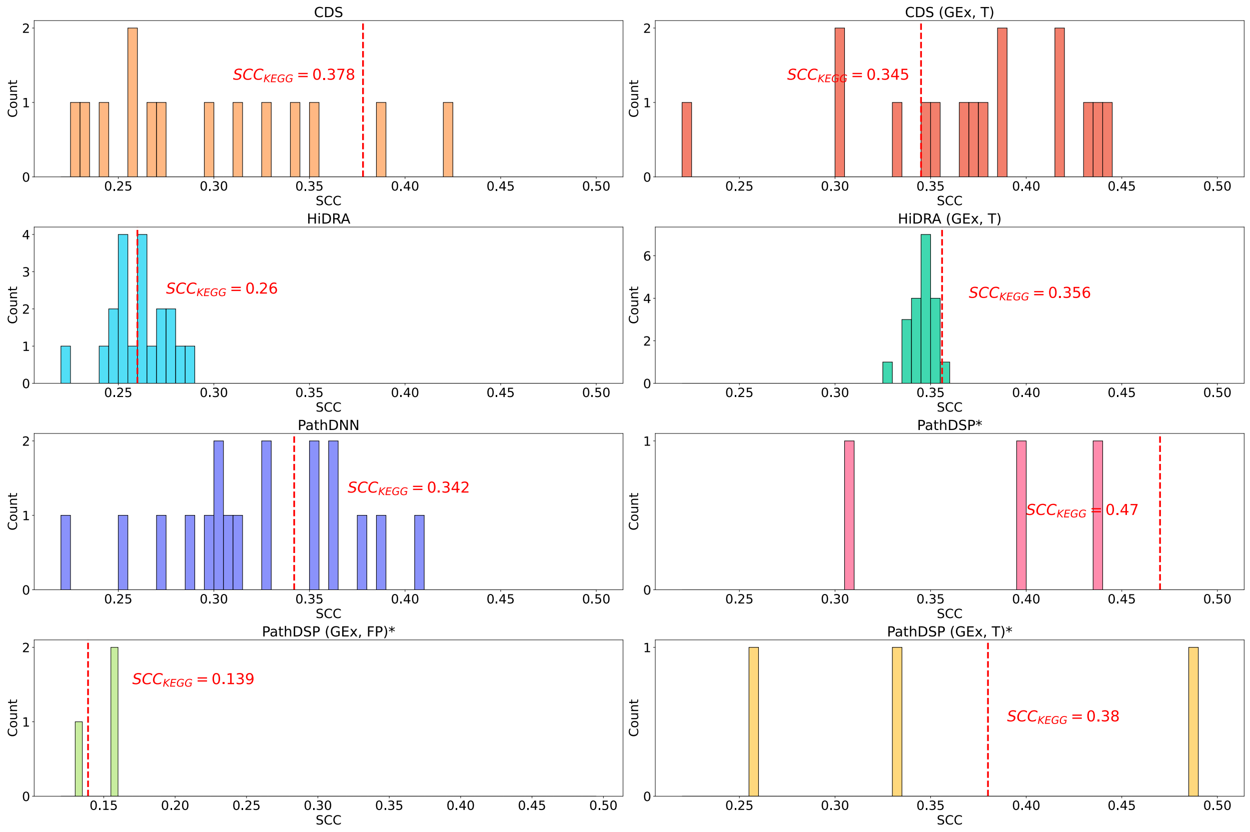


**Figure S7:** Performance of pathway-based models using KEGG or randomly generated pathways. The histograms show the distribution of mean SCC of random pathway baselines using the LDO validation scheme. Vertical dashed red lines show SCC of the model when using KEGG pathways. In this figure, CDS corresponds to CDS (GEx, CNV, T), HiDRA corresponds to HiDRA (GEx, FP), PathDNN corresponds to PathDNN (GEx, T), and PathDSP corresponds to PathDSP (GEx, CNV, MuT, FP, T) in Table 2. *Since PathDSP requires 1000 permutation tests for each type of input data, only three random pathway baselines were constructed due to its extremely high computational requirement.

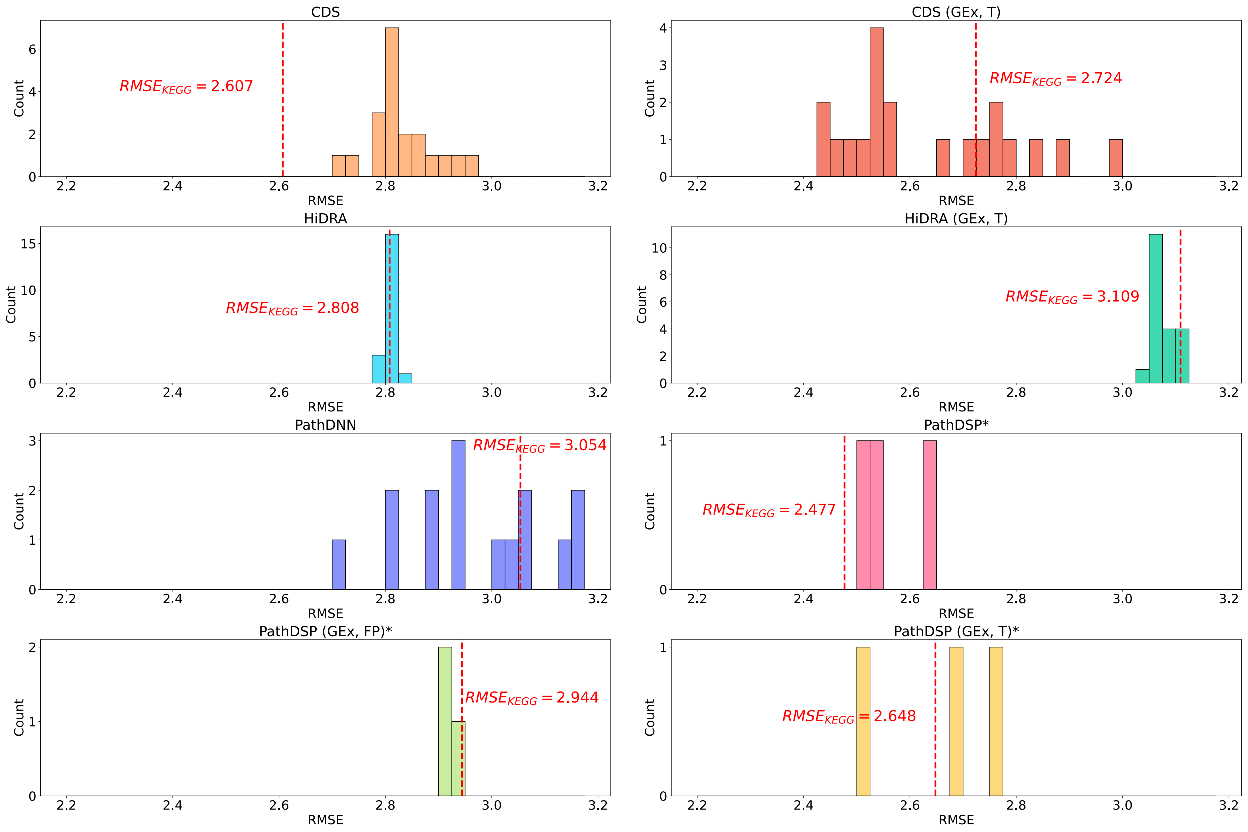


**Figure S8:** Performance of pathway-based models using KEGG or randomly generated pathways. The histograms show the distribution of mean RMSE of random pathway baselines using the LDO validation scheme. Vertical dashed red lines show RMSE of the model when using KEGG pathways. In this figure, CDS corresponds to CDS (GEx, CNV, T), HiDRA corresponds to HiDRA (GEx, FP), PathDNN corresponds to PathDNN (GEx, T), and PathDSP corresponds to PathDSP (GEx, CNV, MuT, FP, T) in Table 2. *Since PathDSP requires 1000 permutation tests for each type of input data, only three random pathway baselines were constructed due to its extremely high computational requirement.

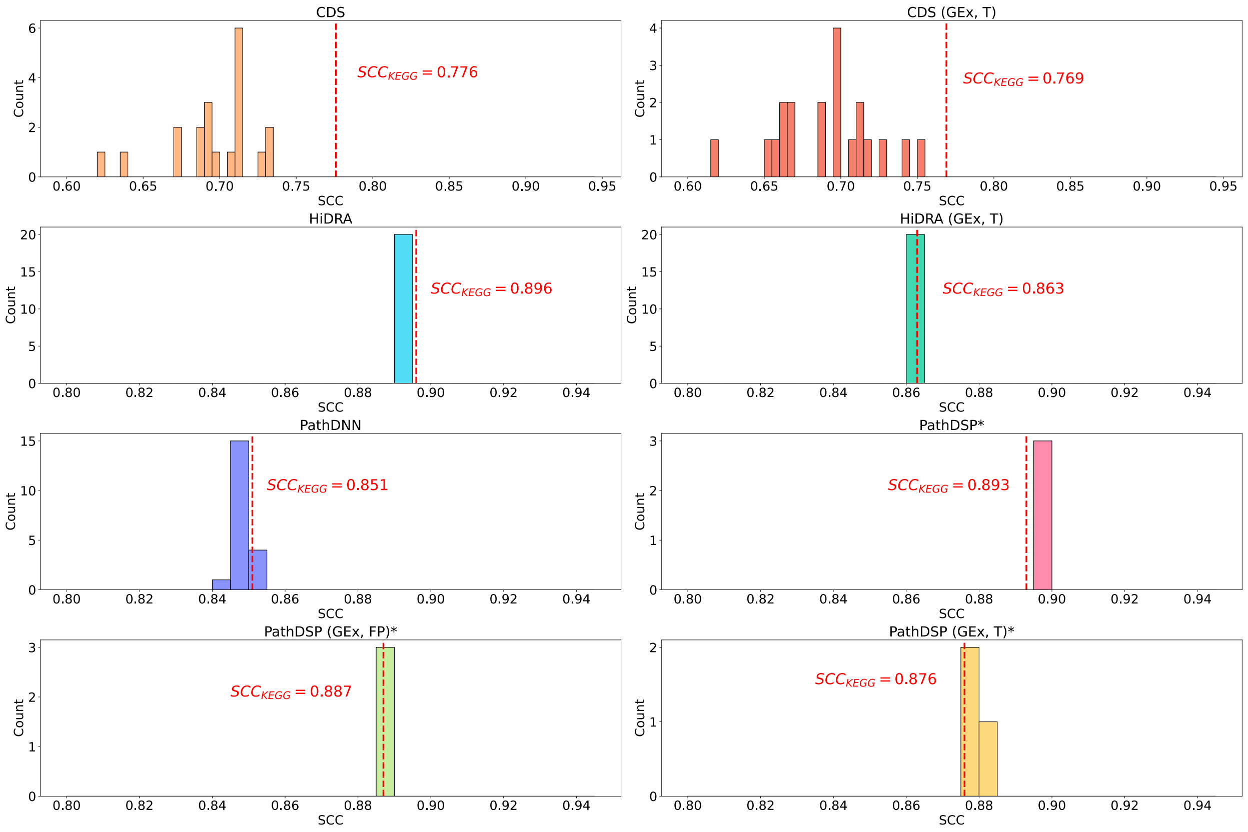


**Figure S9:** Performance of pathway-based models using KEGG or randomly generated pathways. The histograms show the distribution of mean SCC of random pathway baselines using the LPO validation scheme. Vertical dashed red lines show SCC of the model when using KEGG pathways. In this figure, CDS corresponds to CDS (GEx, CNV, T), HiDRA corresponds to HiDRA (GEx, FP), PathDNN corresponds to PathDNN (GEx, T), and PathDSP corresponds to PathDSP (GEx, CNV, MuT, FP, T) in Table 2. *Since PathDSP requires 1000 permutation tests for each type of input data, only three random pathway baselines were constructed due to its extremely high computational requirement.

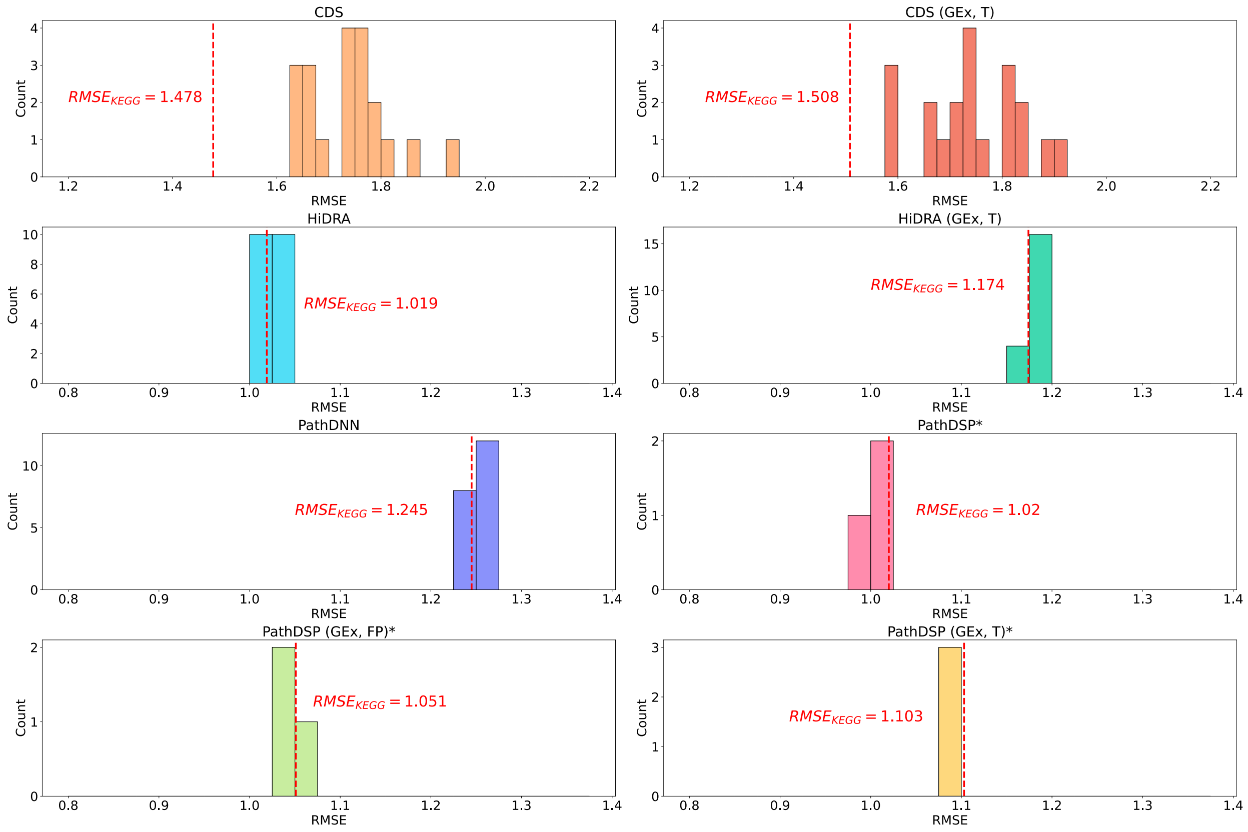


**Figure S10:** Performance of pathway-based models using KEGG or randomly generated pathways. The histograms show the distribution of mean RMSE of random pathway baselines using the LPO validation scheme. Vertical dashed red lines show RMSE of the model when using KEGG pathways. In this figure, CDS corresponds to CDS (GEx, CNV, T), HiDRA corresponds to HiDRA (GEx, FP), PathDNN corresponds to PathDNN (GEx, T), and PathDSP corresponds to PathDSP (GEx, CNV, MuT, FP, T) in Table 2. *Since PathDSP requires 1000 permutation tests for each type of input data, only three random pathway baselines were constructed due to its extremely high computational requirement.

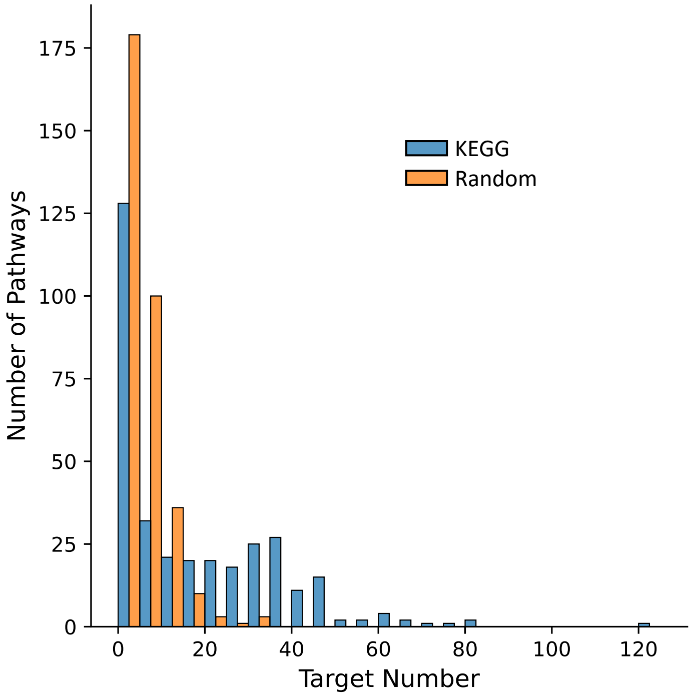


**Figure S11:** Distribution of pathway membership for drug targets in KEGG pathways (blue) and random pathways (orange). The member genes of each biological pathway were examined and any drug target gene among these member genes were identified. Similarly, for randomly generated pathways, the procedure was repeated. Since multiple random pathway collections were generated, the mean of drug targets for each pathway was used in the figure above (orange bars).

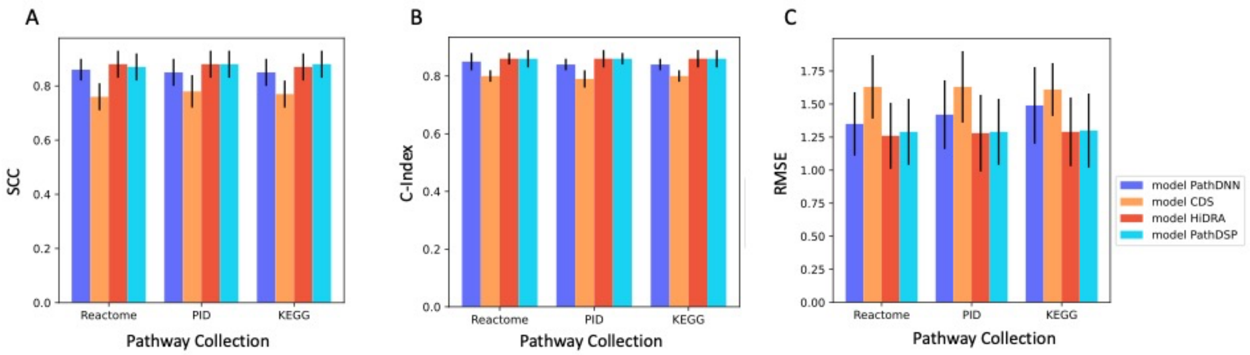


**Figure S12:** Performance of pathway-based models using different pathway collections. Mean and standard deviation are calculated across cell lines using the LCO evaluation. Standard deviations are shown as error bars. A) Spearman Correlation Coefficient (SCC), B) Concordance Index (C-Index), C) Root Mean Squared Error (RMSE). This figure corresponds to Table 5 of the main manuscript.

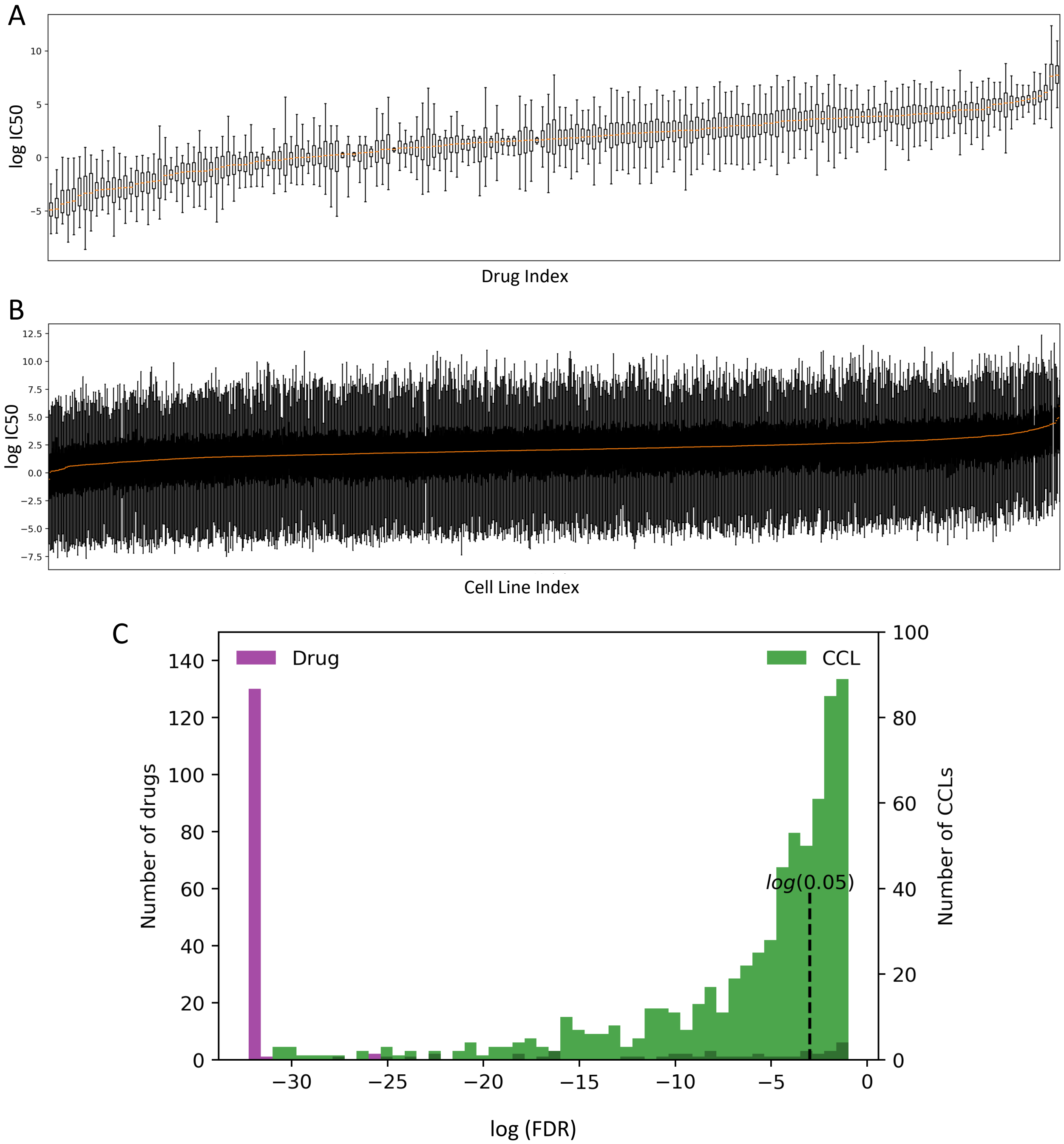


**Figure S13:** Drug and cell line specificity of log IC50 values in our dataset. A) Each box plot shows the distribution of log IC50 values of a single drug (x-axis) across all cancer cell lines (CCLs). B) Each box plot shows the distribution of log IC50 values of a single CCL (x-axis) across all drugs. C) The histogram of false discovery rates (FDRs) obtained from comparing the local distribution of log IC50 values per drug (purple) or per CCL (green) and the global distribution of the log IC50 values using Mann-Whitney U tests. The x-axis shows the log (FDR) values and the dashed vertical line shows the log (0.05) threshold. FDR values are obtained by correcting the p-values for multiple tests using Benjamini-Hochberg method.
