## Supplementary File S2 for "Interpretable deep learning architectures for improving drug response prediction performance: myth or reality?"

**A guideline for architecture, hyperparameter tuning, and training of models**

**Hyperparameter Tuning and Model Training**

We split our data into five folds, where the ratio between train, validation, and test set is 3:1:1. For each model with each pathway collection (KEGG [1], PID [2], Reactome [3]) under each validation scheme (LPO, LCO, LDO), we used the validation set and Ray Tune [4] to perform hyperparameter tuning. Therefore, a total of 9 hyperparameter tuning processes (3 pathway collections $\times$ 3 validation schemes) were done for each model. For each model, the hyperparameter search grid is the same for the three validation schemes, but is different for the different pathway collections, since the number of pathways determines the size of network layers and varies between different pathway collections. Early stopping was also applied to determine the most optimal training epoch and the maximum training epoch is set to 100. Mini-batch gradient descent was applied to all models. The model-specific random pathway baselines used the same set of optimal hyperparameters as their corresponding pathway-based models. All models are implemented using the PyTorch [5] framework and the pathway-incorporated models are re-implemented following the architecture and pipeline described in the original studies.

For each model with each pathway collection, we first show the hyperparameter search grid. Then, the final selection of optimal hyperparameters for each validation scheme are listed in the same table, where the listing order is LPO, LCO, LDO (from left to right). If only one value is listed for a particular hyperparameter, it means that the model used the same hyperparameter value under all three validation schemes.

**MLP (GEx, FP)**

The MLP (GEx, FP) model is a five-layer MLP that uses CCL gene expression profiles as CCL features and Morgan fingerprints as drug features.

***MLP (GEx, FP) – KEGG***

| Hyperparameter | Search Grid |
|---|---|
| 1st hidden layer | [2048, 1024] |
| 2nd hidden layer | [1024, 512] |
| 3rd hidden layer | [512, 256] |
| 4th hidden layer | [128, 64] |
| Learning rate | [1e-3, 1e-4, 1e-5] |
| Weight decay | [1e-5, 0] |

| Hyperparameter | Optimal Hyperparameter Value (LPO, LCO, LDO) |
|---|---|
| Hidden layers | [2048, 1024, 512, 128], [2048, 1024, 256, 64], [2048, 1024, 256, 64] |
| Learning rate | [1e-4], [1e-4], [1e-5] |
| Weight decay | [0], [1e-5], [1e-5] |
| Activation function | ReLU |
| Optimization function | Adam |
| Epoch | [89], [10], [2] |

***MLP (GEx, FP) – PID***

| Hyperparameter | Search Grid |
|---|---|
| 1st hidden layer | [1024, 512] |
| 2nd hidden layer | [512, 256] |
| 3rd hidden layer | [128, 64] |
| 4th hidden layer | [64, 32] |
| Learning rate | [1e-3, 1e-4, 1e-5] |
| Weight decay | [1e-5, 0] |

| Hyperparameter | Value (LPO, LCO, LDO) |
|---|---|
| Hidden layers | [1024, 512, 128, 64], [1024, 256, 128, 64], [1024, 256, 64, 64] |
| Learning rate | [1e-4], [1e-4], [1e-5] |
| Weight decay | 1e-5 |
| Activation function | ReLU |
| Optimization function | Adam |
| Epoch | [100], [11], [4] |

***MLP (GEx, FP) – Reactome***

| Hyperparameter | Search Grid |
|---|---|
| $1^{st}$ hidden layer | [2048, 1024] |
| $2^{nd}$ hidden layer | [1024, 512] |
| $3^{rd}$ hidden layer | [512, 256] |
| $4^{th}$ hidden layer | [256, 128, 64] |
| Learning rate | [1e-3, 1e-4, 1e-5] |
| Weight decay | [1e-5, 0] |

| Hyperparameter | Value (LPO, LCO, LDO) |
|---|---|
| Hidden layers | [2048, 1024, 512, 64], [2048, 512, 512, 256], [1024, 1024, 256, 128] |
| Learning rate | [1e-4], [1e-4], [1e-5] |
| Weight decay | [0], [1e-5], [1e-5] |
| Activation function | ReLU |
| Optimization function | Adam |
| Epoch | [96], [14], [2] |

**MLP (GEx, T)**

***MLP (GEx, T) – KEGG***

| Hyperparameter | Search Grid |
|---|---|
| $1^{st}$ hidden layer | [2048, 1024] |
| $2^{nd}$ hidden layer | [1024, 512] |
| $3^{rd}$ hidden layer | [512, 256] |
| $4^{th}$ hidden layer | [128, 64] |
| Learning rate | [1e-3, 1e-4, 1e-5] |
| Weight decay | [1e-5, 0] |

| Hyperparameter | Value (LPO, LCO, LDO) |
|---|---|
| Hidden layers | [1024, 512, 256, 128], [2048, 512, 256, 128], [1024, 1024, 256, 64] |
| Learning rate | 1e-4 |
| Weight decay | 1e-5 |
| Activation function | ReLU |
| Optimization function | Adam |
| Epoch | [80], [51], [2] |

***MLP (GEx, T) – PID***

| **Hyperparameter** | **Search Grid** |
|---|---|
| 1st hidden layer | [1024, 512] |
| 2nd hidden layer | [512, 256] |
| 3rd hidden layer | [128, 64] |
| 4th hidden layer | [64, 32] |
| Learning rate | [1e-3, 1e-4, 1e-5] |
| Weight decay | [1e-5, 0] |

| **Hyperparameter** | **Value (LPO, LCO, LDO)** |
|---|---|
| Hidden layers | [512, 256, 256, 128], [1024, 256, 256, 64], [512, 512, 256, 64] |
| Learning rate | 1e-4 |
| Weight decay | [1e-5], [1e-5], [0] |
| Activation function | ReLU |
| Optimization function | Adam |
| Epoch | [67], [53], [11] |

***MLP (GEx, T) – Reactome***

| **Hyperparameter** | **Search Grid** |
|---|---|
| 1st hidden layer | [2048, 1024] |
| 2nd hidden layer | [1024, 512] |
| 3rd hidden layer | [512, 256] |
| 4th hidden layer | [256, 128, 64] |
| Learning rate | [1e-3, 1e-4, 1e-5] |
| Weight decay | [1e-5, 0] |

| **Hyperparameter** | **Value (LPO, LCO, LDO)** |
|---|---|
| Hidden layers | [2048, 512, 512, 128], [2048, 1024, 512, 256], [2048, 512, 256, 128] |
| Learning rate | [1e-4], [1e-4], [1e-5] |
| Weight decay | 0 |
| Activation function | ReLU |
| Optimization function | Adam |
| Epoch | [92], [66], [1] |

**PathDNN**

The PathDNN model uses CCL gene expression profile as CCL features and drug target data as drug features. It contains three hidden layers, where the first hidden layer is a pathway layer with each node representing a pathway. Our implementation of PathDNN resembles the original model architecture and follows the original study's choice for optimization function, activation function, and the inclusion of a drop layer. The original study used Adam and stochastic gradient descent (SGD) optimizers, where the Adam optimizer was used first to train the model and the SGD optimizer was used later (for the last 10% of training epochs) when the model has mainly converged. The figure below illustrates the model architecture of PathDNN. The size of the pathway layer is determined by the number of pathways in the pathway collection, therefore it is not tuneable.

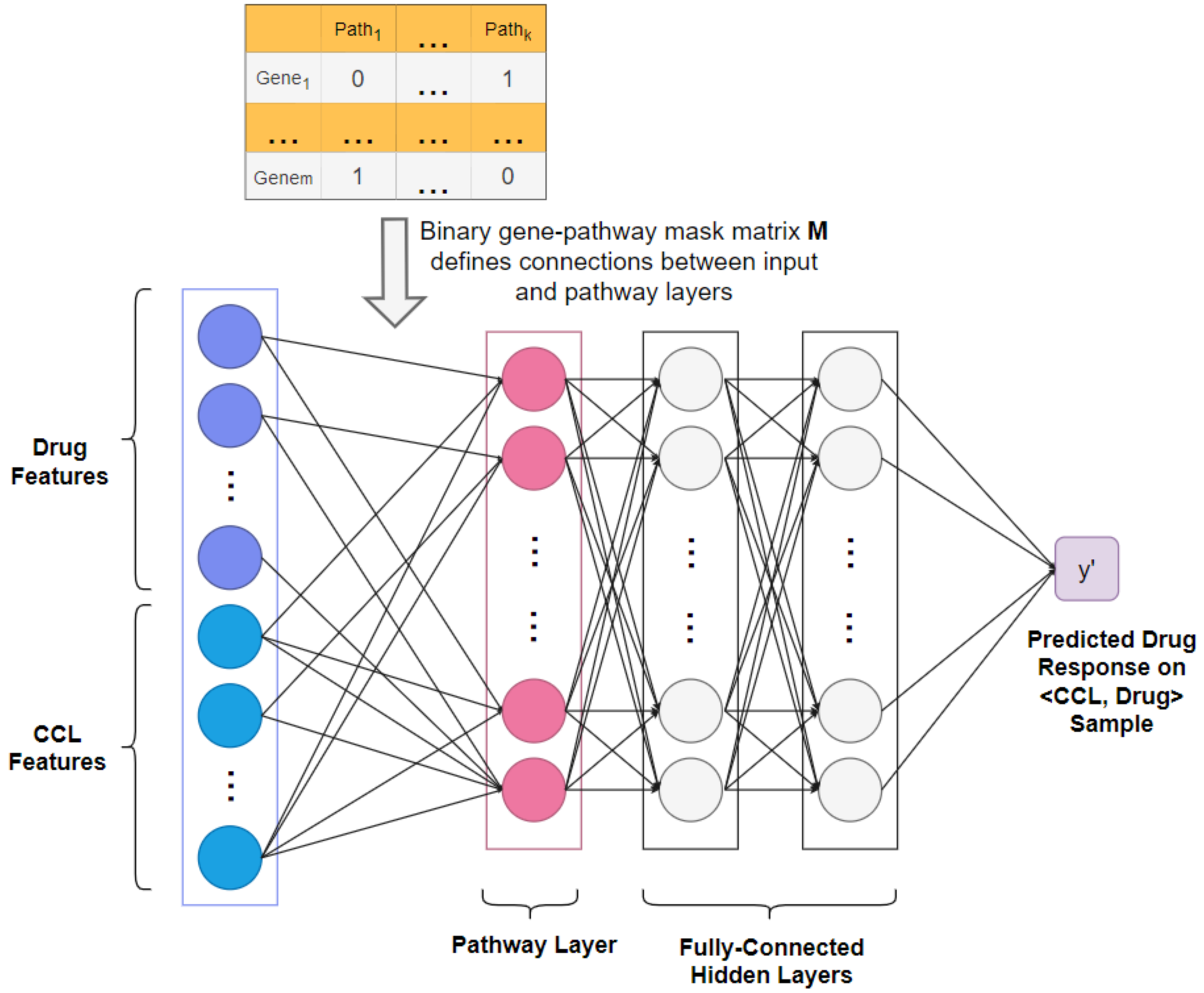

***PathDNN – KEGG***

| Hyperparameter | Search Grid |
|---|---|
| 1st hidden layer | [512, 256, 128] |
| 2nd hidden layer | [256, 128] |
| Drop rate | [0.05, 0.1, 0.2] |
| Learning rate (Adam) | [1e-4, 2e-4] |
| Weight decay (Adam) | [1e-4, 0] |
| Learning rate (SGD) | [1e-5, 5e-5] |
| Weight decay (SGD) | [1e-4, 0] |

| Hyperparameter | Value (LPO, LCO, LDO) |
|---|---|
| Pathway layer | 332 |
| Hidden layers | [256, 256], [512, 256], [256, 256] |
| Learning rate [Adam, SGD] | [2e-4, 1e-5], [2e-4, 1e-5], [2e-4, 5e-5] |
| Weight decay [Adam, SGD] | [0, 1e-4] |
| Drop rate | [0.2], [0.05], [0.2] |
| Activation function | ReLU |
| Optimization function | Adam, stochastic gradient descent (SGD) |
| Epoch | [57], [37], [39] |

***PathDNN – PID***

| Hyperparameter | Search Grid |
|---|---|
| 1st hidden layer | [512, 256, 128] |
| 2nd hidden layer | [256, 128] |
| Drop rate | [0.05, 0.1, 0.2] |
| Learning rate (Adam) | [1e-4, 2e-4] |
| Weight decay (Adam) | [1e-4, 0] |
| Learning rate (SGD) | [1e-5, 5e-5] |
| Weight decay (SGD) | [1e-4, 0] |

| Hyperparameter | Value (LPO, LCO, LDO) |
|---|---|
| Pathway layer | 196 |
| Hidden layers | [512, 256], [128, 128], [256, 256] |
| Learning rate [Adam, SGD] | [1e-4, 5e-5], [2e-4, 1e-5], [2e-4, 1e-5] |
| Weight decay [Adam, SGD] | [0, 0], [0, 1e-4], [0, 0] |
| Drop rate | 0.2 |
| Activation function | ReLU |
| Optimization function | Adam, stochastic gradient descent (SGD) |
| Epoch | [38], [17], [1] |

***PathDNN – Reactome***

| Hyperparameter | Search Grid |
|---|---|
| 1st hidden layer | [2048, 1024, 512] |
| 2nd hidden layer | [512, 256, 128] |
| Drop rate | [0.05, 0.1, 0.2] |
| Learning rate (Adam) | [1e-4, 2e-4] |
| Weight decay (Adam) | [1e-4, 0] |
| Learning rate (SGD) | [1e-5, 5e-5] |
| Weight decay (SGD) | [1e-4, 0] |

| Hyperparameter | Value (LPO, LCO, LDO) |
|---|---|
| Pathway layer | 1608 |
| Hidden layers | [2048, 256], [2048, 512], [512, 128] |
| Learning rate [Adam, SGD] | [2e-4, 1e-5], [2e-4, 5e-5], [1e-4, 5e-5] |
| Weight decay [Adam, SGD] | [1e-4, 1e-4], [1e-4, 0], [0, 1e-4] |
| Drop rate | [0.1], [0.05], [0.05] |
| Activation function | ReLU |
| Optimization function | Adam, stochastic gradient descent (SGD) |
| Epoch | [16], [10], [18] |

**ConsDeepSignaling (CDS)**

ConsDeepSignaling (CDS) takes gene expression profile and copy number variation as CCL features and drug target information as drug features. The three types of input features are concatenated in the input layer. The input layer is followed by a gene layer, where the connections between these two layers are defined by a binary mask matrix $M_{XG}$. Each node in the gene layer is connected to its corresponding input features. The gene layer is followed by a pathway layer, where a binary mask matrix $M_{GP}$ defines the connections between these two layers based on pathway membership. Finally, the pathway layer is followed by three fully-connected hidden layers to complete the prediction network. The size of the gene layer is determined by the number of unique genes in the input data and the size of the pathway layer is determined by the number of pathways in the pathway collection. As a result, the sizes of these two layers are fixed and not tuneable. Our implementation of CDS resembles the original model architecture and follows the original study's choice for optimization function and activation function. The figure below illustrates the model architecture of CDS.

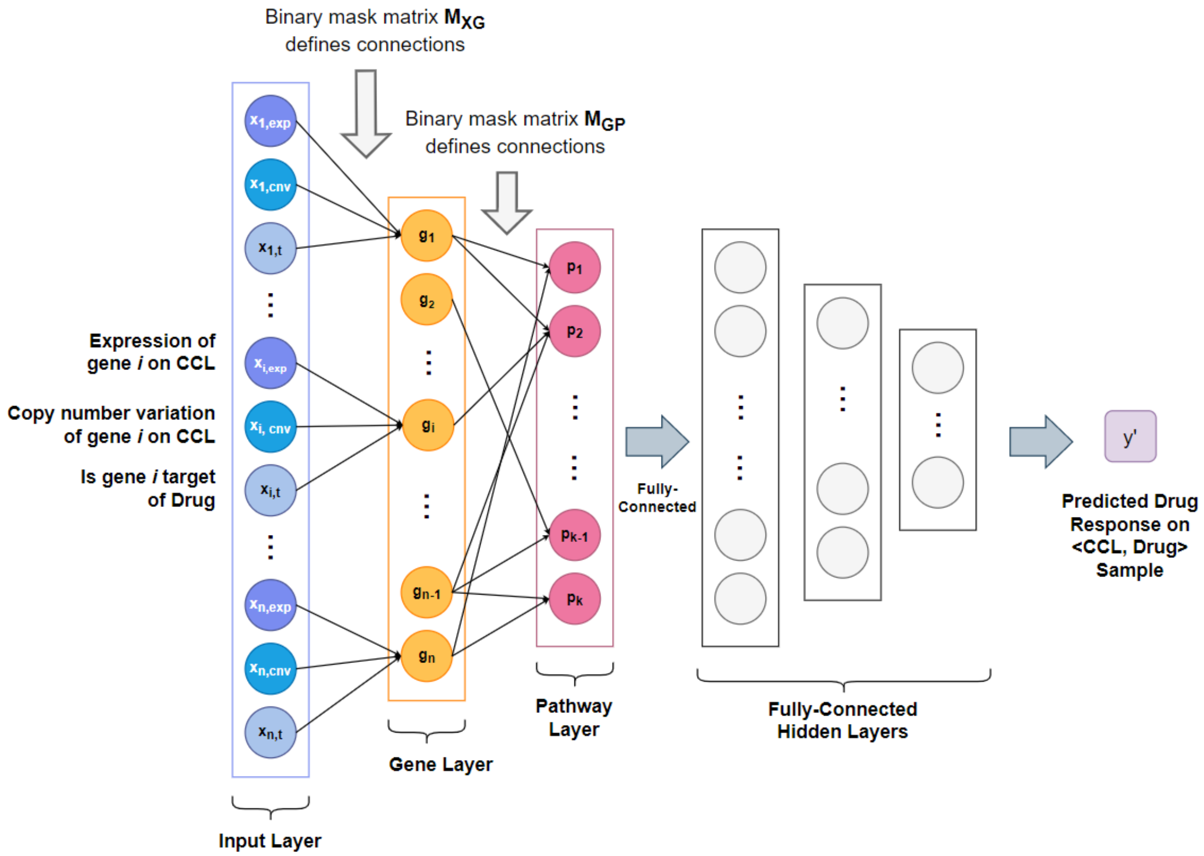

***CDS – KEGG***

| Hyperparameter | Search Grid |
|---|---|
| 1st hidden layer | [512, 256, 128] |
| 2nd hidden layer | [256, 128] |
| 3rd hidden layer | [128, 64] |
| Learning rate | [1e-4, 2e-4, 5e-4, 1e-3] |
| Weight decay | Default* |

* The weight decay of the Adam optimizer was set to default, since the original study did not report a value for the weight decay.

| Hyperparameter | Value (LPO, LCO, LDO) |
|---|---|
| Gene layer | 5511 |
| Pathway layer | 332 |
| Hidden layers | [512, 128, 64], [256, 256, 64], [256, 64, 32] |
| Learning rate | [1e-3], [1e-3], [1e-4] |
| Weight decay | default |
| Activation function | ReLU |
| Optimization function | Adam |
| Epoch | [46], [23], [5] |

***CDS – PID***

| Hyperparameter | Search Grid |
|---|---|
| 1st hidden layer | [512, 256, 128] |
| 2nd hidden layer | [256, 128] |
| 3rd hidden layer | [128, 64] |
| Learning rate | [1e-4, 2e-4, 5e-4, 1e-3] |
| Weight decay | Default* |

* The weight decay of the Adam optimizer was set to default, since the original study did not report a value for the weight decay.

| Hyperparameter | Value (LPO, LCO, LDO) |
|---|---|
| Gene layer | 2078 |
| Pathway layer | 196 |
| Hidden layers | [128, 128, 64], [128, 128, 128], [128, 128, 128] |
| Learning rate | 1e-3 |
| Weight decay | default |
| Activation function | ReLU |
| Optimization function | Adam |
| Epoch | [50], [79], [5] |

***CDS – Reactome***

| Hyperparameter | Search Grid |
|---|---|
| 1st hidden layer | [1024, 512] |
| 2nd hidden layer | [512, 256] |
| 3rd hidden layer | [256, 128, 64] |
| Learning rate | [1e-4, 2e-4, 5e-4, 1e-3] |
| Weight decay | default* |

* The weight decay of the Adam optimizer was set to default, since the original study did not report a value for the weight decay.

| Hyperparameter | Value (LPO, LCO, LDO) |
|---|---|
| Gene layer | 7831 |
| Pathway layer | 1608 |
| Hidden layers | [1024, 256, 128], [1024, 512, 64], [1024, 256, 128] |
| Learning rate | [1e-3], [1e-3], [2e-4] |
| Weight decay | default |
| Activation function | ReLU |
| Optimization function | Adam |
| Epoch | [39], [32], [1] |

**CDS (GEx, T)**

The CDS (GEx, T) has the same model architecture and pipeline as the original CDS model, except the input data only has two types of input features (gene expression profiles, drug target information), as opposed to three (gene expression profiles, copy number variation, drug target information). The hyperparameter search grid for CDS (GEx, T) is the same as CDS.

***CDS (GEx, T) – KEGG***

| **Hyperparameter** | **Value (LPO, LCO, LDO)** |
|---|---|
| Gene layer | 5511 |
| Pathway layer | 332 |
| Hidden layers | [128, 256, 64], [512, 128, 128], [512, 256, 64] |
| Learning rate | [1e-3], [5e-4], [2e-4] |
| Weight decay | default |
| Activation function | ReLU |
| Optimization function | Adam |
| Epoch | [84], [46], [5] |

***CDS (GEx, T) – PID***

| **Hyperparameter** | **Value (LPO, LCO, LDO)** |
|---|---|
| Gene layer | 2078 |
| Pathway layer | 196 |
| Hidden layers | [512, 256, 128], [128, 128, 128], [256, 256, 64] |
| Learning rate | [1e-3], [5e-4], [5e-4] |
| Weight decay | default |
| Activation function | ReLU |
| Optimization function | Adam |
| Epoch | [78], [64], [1] |

***CDS (GEx, T) – Reactome***

| **Hyperparameter** | **Value (LPO, LCO, LDO)** |
|---|---|
| Gene layer | 7831 |
| Pathway layer | 1608 |
| Hidden layers | [1024, 128, 64], [1024, 128, 128], [512, 256, 256] |
| Learning rate | [5e-4], [1e-3], [5e-4] |
| Weight decay | default |
| Activation function | ReLU |
| Optimization function | Adam |
| Epoch | [36], [43], [5] |

**HiDRA**

HiDRA has a hierarchical network architecture. The figure below illustrates the model architecture. The drug attention network consists of two fully-connected hidden layers and takes in drug features (drug target information) to generate new encoding of the drug features. These drug encodings are then concatenated with gene expression data of pathway genes in the gene-level network to generate pathway activation scores. The gene-level network contains multiple smaller networks, where each smaller network is dedicated to a pathway and the output of the smaller network is the pathway activation score. The output of the gene-level network is then concatenated with the output of the drug attention network to act as the input to the pathway-level network. The output of the pathway-level network is weighted pathway activation score and is then concatenated with the output of the drug network to generate the final input to the drug response prediction network. The reimplementation of HiDRA follows the exact same architecture and pipeline as the original study. The choices for optimizer and activation function also follow that of the original study.

Since HiDRA contains multiple network modules (the gene-level network contains 332 smaller networks for KEGG, 196 for PID, 1608 for Reactome), the computation time and the number of tuneable hyperparameters are significantly higher compared to the other pathway-based models. Due to computation constraints, we did not perform hyperparameter tuning on the hidden layer size of the drug network and the response network. We just used the reported drug network and response network hidden layer size of the original study. We decided to do this because the size of our drug features is the same as the original study of HiDRA (both used 512-bit Morgan fingerprints). Moreover, the response network only contains one hidden layer and the input data size to the response network of our study is similar to that of the original study of HiDRA. Due to the above reasons, the reported hidden layer size for these two sub-networks in the original study were used in our reimplementation of HiDRA.

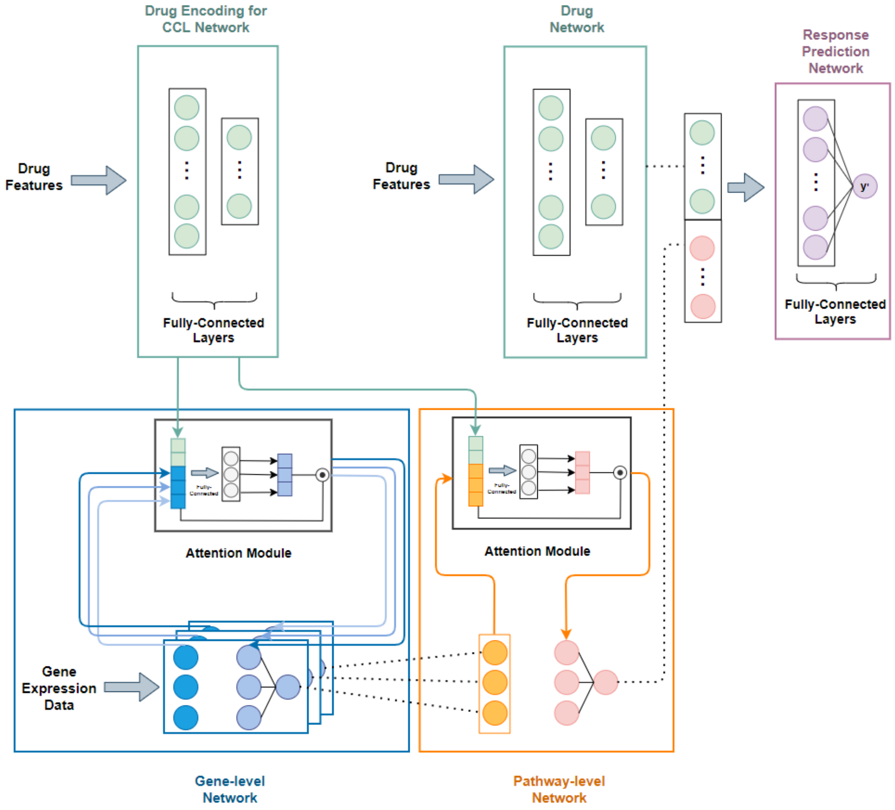

***HiDRA – KEGG***

| **Hyperparameter** | **Search Grid** |
|---|---|
| Drug encoding for CCL network layer 1 | [256, 128] |
| Drug encoding for CCL network layer 2 | [64, 32] |
| Encoding for gene-level network | $[\frac{n_i}{4}, \frac{n_i}{8}, \frac{n_i}{16}]$* |
| Encoding for pathway-level network | $[\frac{n_i}{4}, \frac{n_i}{8}, \frac{n_i}{16}]$* |
| Learning rate | [1e-4, 2e-4, 5e-4, 1e-3] |

* $n_i$ is the number of member genes in pathway i. The gene-level network contains multiple smaller networks, where each smaller network is dedicated to calculating pathway activation score for a single pathway. Since the number of pathway member genes varies, the size of the hidden layer within the smaller network is proportional to $n_i$.

| **Hyperparameter** | **Value (LPO, LCO, LDO)** |
|---|---|
| Drug network layers | [256, 128] |
| Drug attention network layers | [256, 64] |
| Drug encoding for gene-level network | $[\frac{n_i}{4}]$ |
| Drug encoding for pathway-level network | $[\frac{n_i}{16}]$, $[\frac{n_i}{16}]$, $[\frac{n_i}{8}]$ |
| Drug response network | 128 |
| Learning rate | [1e-3], [2e-4], [1e-4] |
| Weight decay | default* |
| Optimization function | Adam |
| Activation function | ReLU |
| Epoch | [40], [25], [25] |

* The weight decay of the Adam optimizer was set to default, since the original study did not report a value for the weight decay.

***HiDRA – PID***

| **Hyperparameter** | **Search Grid** |
|---|---|
| Drug encoding for CCL network layer 1 | [256, 128] |
| Drug encoding for CCL network layer 2 | [64, 32] |
| Encoding for gene-level network | $[\frac{n_i}{4}, \frac{n_i}{8}, \frac{n_i}{16}]$* |
| Encoding for pathway-level network | $[\frac{n_i}{4}, \frac{n_i}{8}, \frac{n_i}{16}]$* |
| Learning rate | [1e-4, 2e-4, 5e-4, 1e-3] |

* $n_i$ is the number of member genes in pathway i. The gene-level network contains multiple smaller networks, where each smaller network is dedicated to calculating pathway activation score for a single pathway. Since the number of pathway member genes varies, the size of the hidden layer within the smaller network is proportional to $n_i$.

| Hyperparameter | Value (LPO, LCO, LDO) |
|---|---|
| Drug network layers | [256, 128] |
| Drug attention network layers | [256, 32], [128, 64], [128, 32] |
| Drug encoding for gene-level network | [$\frac{n_i}{4}$], [$\frac{n_i}{16}$], [$\frac{n_i}{4}$] |
| Drug encoding for pathway-level network | [$\frac{n_i}{8}$], [$\frac{n_i}{8}$], [$\frac{n_i}{16}$] |
| Drug response network | 128 |
| Learning rate | [1e-3] |
| Weight decay | default* |
| Optimization function | Adam |
| Activation function | ReLU |
| Epoch | [12], [6], [1] |

* The weight decay of the Adam optimizer was set to default, since the original study did not report a value for the weight decay.

***HiDRA – Reactome***

| Hyperparameter | Search Grid |
|---|---|
| Drug encoding for CCL network layer 1 | [256, 128] |
| Drug encoding for CCL network layer 2 | [64, 32] |
| Encoding for gene-level network | [$\frac{n_i}{4}$, $\frac{n_i}{8}$, $\frac{n_i}{16}$]* |
| Encoding for pathway-level network | [$\frac{n_i}{4}$, $\frac{n_i}{8}$, $\frac{n_i}{16}$]* |
| Learning rate | [1e-4, 2e-4, 5e-4, 1e-3] |

* $n_i$ is the number of member genes in pathway i. The gene-level network contains multiple smaller networks, where each smaller network is dedicated to calculating pathway activation score for a single pathway. Since the number of pathway member genes varies, the size of the hidden layer within the smaller network is proportional to $n_i$.

| Hyperparameter | Value (LPO, LCO, LDO) |
|---|---|
| Drug network layers | [256, 128] |
| Drug attention network layers | [256, 64], [256, 64], [128, 64] |
| Drug encoding for gene-level network | [$\frac{n_i}{4}$] |
| Drug encoding for pathway-level network | [$\frac{n_i}{8}$], [$\frac{n_i}{16}$], [$\frac{n_i}{8}$] |
| Drug response network | 128 |
| Learning rate | [1e-3], [2e-4], [2e-4] |
| Weight decay | default* |
| Optimization function | Adam |
| Activation function | ReLU |
| Epoch | [38], [4], [1] |

* The weight decay of the Adam optimizer was set to default, since the original study did not report a value for the weight decay.

**HiDRA (GEx, T)**

HiDRA (GEx, T) has the exact same architecture and pipeline as HiDRA, except it uses drug target information as drug features, as opposed to Morgan fingerprints used by HiDRA. The hyperparameter search grid for HiDRA (GEx, T) is the same as HiDRA.

***HiDRA (GEx, T) – KEGG***

| **Hyperparameter** | **Value (LPO, LCO, LDO)** |
|---|---|
| Drug network layers | [256, 128] |
| Drug attention network layers | [128, 32], [256, 64], [256, 32] |
| Drug encoding for gene-level network | $[\frac{n_i}{4}]$, $[\frac{n_i}{8}]$, $[\frac{n_i}{16}]$ |
| Drug encoding for pathway-level network | $[\frac{n_i}{16}]$, $[\frac{n_i}{16}]$, $[\frac{n_i}{8}]$ |
| Drug response network | 128 |
| Learning rate | [1e-3], [1e-3], [1e-4] |
| Weight decay | Default* |
| Optimization function | Adam |
| Activation function | ReLU |
| Epoch | [7], [6], [1] |

* The weight decay of the Adam optimizer was set to default, since the original study did not report a value for the weight decay.

***HiDRA (GEx, T) – PID***

| **Hyperparameter** | **Value (LPO, LCO, LDO)** |
|---|---|
| Drug network layers | [256, 128] |
| Drug attention network layers | [128, 64], [256, 32], [256, 64] |
| Drug encoding for gene-level network | $[\frac{n_i}{4}]$, $[\frac{n_i}{16}]$, $[\frac{n_i}{8}]$ |
| Drug encoding for pathway-level network | $[\frac{n_i}{8}]$, $[\frac{n_i}{8}]$, $[\frac{n_i}{16}]$ |
| Drug response network | 128 |
| Learning rate | [1e-3], [1e-3], [5e-4] |
| Weight decay | default* |
| Optimization function | Adam |
| Activation function | ReLU |
| Epoch | [6], [4], [11] |

* The weight decay of the Adam optimizer was set to default, since the original study did not report a value for the weight decay.

***HiDRA (GEx, T) – Reactome***

| **Hyperparameter** | **Value (LPO, LCO, LDO)** |
|---|---|
| Drug network layers | [256, 128] |
| Drug attention network layers | [256, 64], [128, 32], [128, 32] |
| Drug encoding for gene-level network | [$\frac{n_i}{4}$], [$\frac{n_i}{4}$], [$\frac{n_i}{16}$] |
| Drug encoding for pathway-level network | [$\frac{n_i}{16}$], [$\frac{n_i}{16}$], [$\frac{n_i}{8}$] |
| Drug response network | 128 |
| Learning rate | [1e-3], [5e-4], [2e-4] |
| Weight decay | default* |
| Optimization function | Adam |
| Activation function | ReLU |
| Epoch | [8], [4], [3] |

* The weight decay of the Adam optimizer was set to default, since the original study did not report a value for the weight decay.

**PathDSP**

The architecture of PathDSP adopts that of a five-layer MLP. The drug target, gene expression, somatic mutation, and copy number variation data are processed using pathway enrichment analysis to form matrices of enrichment scores, which act as input features to the model. The figure below illustrates the model architecture. The enrichment score matrices are organized with columns representing pathways and rows representing drugs or CCLs. For each training sample of <CCL, Drug> pair, the corresponding Morgan fingerprints and pathway enrichment scores are concatenated to form the input feature vector. Our implementation of PathDSP resembles the original model architecture and follows the original study's choice for optimizer, activation function, and the inclusion of a drop layer.

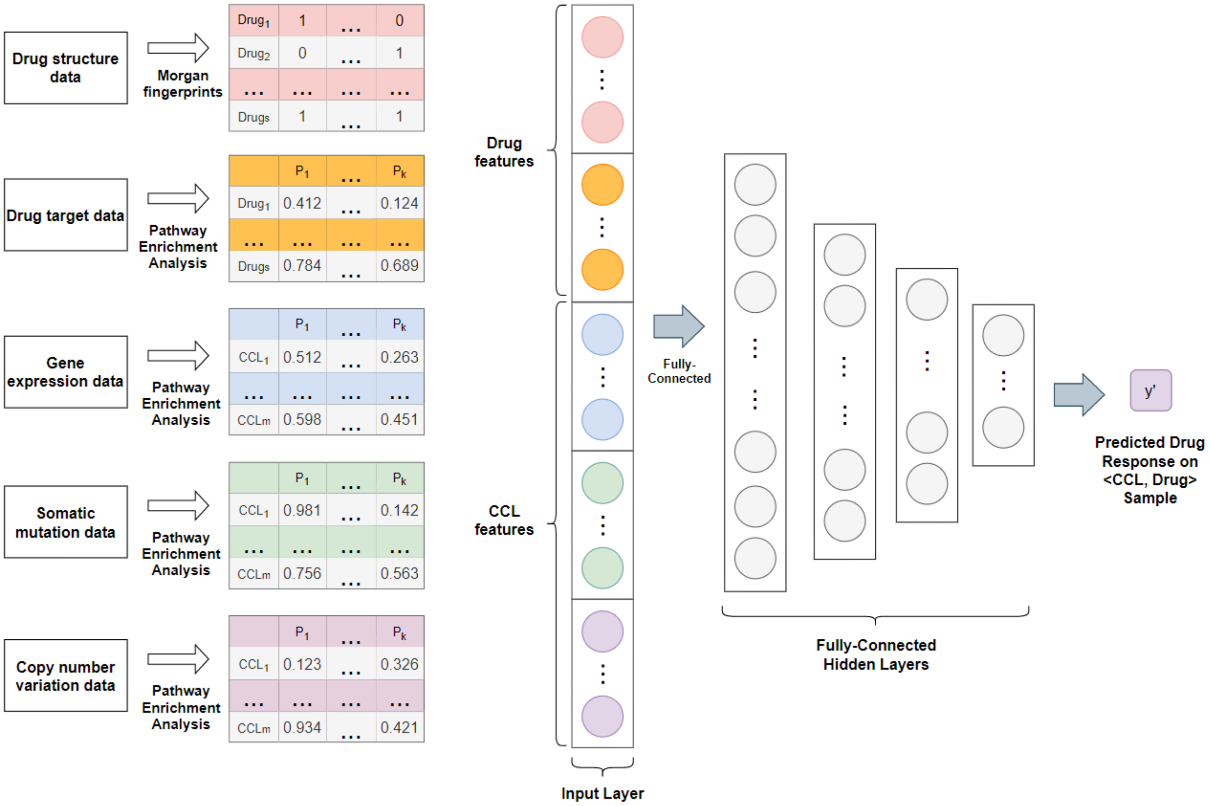

***PathDSP – KEGG***

| Hyperparameter | Search Grid |
|---|---|
| $1^{st}$ hidden layer | [1024, 512] |
| $2^{nd}$ hidden layer | [512, 256] |
| $3^{rd}$ hidden layer | [256, 128] |
| $4^{th}$ hidden layer | [128, 64] |
| Drop rate | [0.05, 0.1, 0.2] |
| Learning rate | [1e-5, 1e-4, 5e-4, 1e-3] |
| Weight decay | default* |

* The weight decay of the Ada Max optimizer was set to default, since the original study did not report a value for the weight decay.

| Hyperparameter | Value (LPO, LCO, LDO) |
|---|---|
| Hidden layers | [1024, 512, 256, 64], [1024, 256, 256, 64], [512, 256, 128, 128] |
| Learning rate | [1e-3], [1e-3], [1e-4] |
| Drop rate | 0.1 |
| Weight decay | default |
| Optimization function | Ada Max |
| Activation function | ELU |
| Epoch | [99], [7], [24] |

***PathDSP – PID***

| Hyperparameter | Search Grid |
|---|---|
| $1^{st}$ hidden layer | [800, 512] |
| $2^{nd}$ hidden layer | [512, 256] |
| $3^{rd}$ hidden layer | [256, 128] |
| $4^{th}$ hidden layer | [128, 64] |
| Drop rate | [0.05, 0.1, 0.2] |
| Learning rate | [1e-5, 1e-4, 5e-4, 1e-3] |
| Weight decay | default* |

* The weight decay of the Ada Max optimizer is not tuned because the original study did not report a value for the weight decay, therefore a default value of 0 was used.

| Hyperparameter | Value (LPO, LCO, LDO) |
|---|---|
| Hidden layers | [800, 512, 256, 128], [512, 256, 256, 64], [512, 256, 256, 64] |
| Learning rate | [1e-3], [1e-3], [1e-4] |
| Weight decay | default |
| Drop rate | 0.1 |
| Optimization function | Ada Max |
| Activation function | ELU |
| Epoch | [96], [8], [4] |

***PathDSP – Reactome***

| Hyperparameter | Search Grid |
|---|---|
| 1$^{st}$ hidden layer | [3072, 2048] |
| 2$^{nd}$ hidden layer | [2048, 1024] |
| 3$^{rd}$ hidden layer | [800, 512] |
| 4$^{th}$ hidden layer | [512, 256, 128] |
| Drop rate | [0.05, 0.1, 0.2] |
| Learning rate | [1e-5, 1e-4, 5e-4, 1e-3] |
| Weight decay | default* |

* The weight decay of the Ada Max optimizer was set to default, since the original study did not report a value for the weight decay.

| Hyperparameter | Value (LPO, LCO, LDO) |
|---|---|
| Hidden layers | [2048, 2048, 800, 512], [3072, 2048, 512, 256], [2048, 2048, 512, 128] |
| Learning rate | [5e-4], [1e-4], [1e-4] |
| Weight decay | default |
| Drop out | 0.1 |
| Optimization function | Ada Max |
| Activation function | ELU |
| Epoch | [93], [15], [17] |

**PathDSP (GEx, FP)**

The architecture of PathDSP (GEx, FP) is the same as PathDSP, except that the input features only includes gene expression profiles for CCL features and Morgan fingerprints for drug features.

***PathDSP (GEx, FP) – KEGG***

| **Hyperparameter** | **Search Grid** |
|---|---|
| 1st hidden layer | [600, 512] |
| 2nd hidden layer | [512, 256] |
| 3rd hidden layer | [256, 128] |
| 4th hidden layer | [128, 64] |
| Drop rate | [0.05, 0.1, 0.2] |
| Learning rate | [1e-5, 1e-4, 5e-4, 1e-3] |
| Weight decay | default* |

* The weight decay of the Ada Max optimizer was set to default, since the original study did not report a value for the weight decay.

| **Hyperparameter** | **Value (LPO, LCO, LDO)** |
|---|---|
| Hidden layers | [600, 512, 256, 128], [600, 256, 128, 64], [600, 512, 256, 128] |
| Learning rate | [1e-4], [1e-5], [1e-4] |
| Drop out | [0.05], [0.1], [0.1] |
| Optimization function | Ada Max |
| Activation function | ELU |
| Epoch | [69], [15], [13] |

***PathDSP (GEx, FP) – PID***

| **Hyperparameter** | **Search Grid** |
|---|---|
| 1st hidden layer | [800, 512] |
| 2nd hidden layer | [512, 256] |
| 3rd hidden layer | [256, 128] |
| 4th hidden layer | [128, 64] |
| Drop rate | [0.05, 0.1, 0.2] |
| Learning rate | [1e-5, 1e-4, 5e-4, 1e-3] |
| Weight decay | default* |

* The weight decay of the Ada Max optimizer was set to default, since the original study did not report a value for the weight decay.

| Hyperparameter | Value (LPO, LCO, LDO) |
|---|---|
| Hidden layers | [800, 512, 256, 64], [512, 512, 128, 64], [800, 256, 256, 128] |
| Learning rate | [5e-4], [1e-4], [1e-5] |
| Drop out | 0.1 |
| Optimization function | Ada Max |
| Activation function | ELU |
| Epoch | [60], [30], [3] |

***PathDSP (GEx, FP) – Reactome***

| Hyperparameter | Search Grid |
|---|---|
| 1st hidden layer | [1024, 800] |
| 2nd hidden layer | [800, 512] |
| 3rd hidden layer | [512, 256] |
| 4th hidden layer | [256, 128] |
| Drop rate | [0.05, 0.1, 0.2] |
| Learning rate | [1e-5, 1e-4, 5e-4, 1e-3] |
| Weight decay | default* |

* The weight decay of the Ada Max optimizer was set to default, since the original study did not report a value for the weight decay.

| Hyperparameter | Value (LPO, LCO, LDO) |
|---|---|
| Hidden layers | [1024, 512, 512, 256], [800, 800, 512, 256], [1024, 800, 256, 128] |
| Learning rate | [5e-4], [5e-4], [1e-5] |
| Weight decay | default |
| Drop out | 0.1 |
| Optimization function | Ada Max |
| Activation function | ELU |
| Epoch | [81], [20], [12] |

**PathDSP (GEx, T)**
PathDSP (GEx, T) has the same architecture and pipeline as PathDSP, except that the input data only include gene expression profiles and drug target information.

***PathDSP (GEx, T) – KEGG***

| **Hyperparameter** | **Search Grid** |
|---|---|
| 1st hidden layer | [600, 512] |
| 2nd hidden layer | [512, 256] |
| 3rd hidden layer | [256, 128] |
| 4th hidden layer | [128, 64] |
| Drop rate | [0.05, 0.1, 0.2] |
| Learning rate | [1e-5, 1e-4, 5e-4, 1e-3] |
| Weight decay | default* |

* The weight decay of the Ada Max optimizer was set to default, since the original study did not report a value for the weight decay.

| **Hyperparameter** | **Value (LPO, LCO, LDO)** |
|---|---|
| Hidden layers | [512, 512, 256, 128], [600, 512, 256, 64], [600, 256, 128, 64] |
| Learning rate | [1e-4], [1e-4], [1e-5] |
| Weight decay | default |
| Drop out | [0.05], [0.1], [0.1] |
| Optimization function | Ada Max |
| Activation function | ELU |
| Epoch | [70], [29], [18] |

***PathDSP (GEx, T) – PID***

| **Hyperparameter** | **Search Grid** |
|---|---|
| 1st hidden layer | [512, 256] |
| 2nd hidden layer | [256, 128] |
| 3rd hidden layer | [128, 64] |
| 4th hidden layer | [64, 32] |
| Drop rate | [0.05, 0.1, 0.2] |
| Learning rate | [1e-5, 1e-4, 5e-4, 1e-3] |
| Weight decay | default* |

* The weight decay of the Ada Max optimizer was set to default, since the original study did not report a value for the weight decay.

| Hyperparameter | Value (LPO, LCO, LDO) |
|---|---|
| Hidden layers | [512, 256, 128, 64], [512, 256, 128, 64], [256, 128, 128, 32] |
| Learning rate | [1e-3], [5e-4], [1e-5] |
| Weight decay | default |
| Drop out | 0.1 |
| Optimization function | Ada Max |
| Activation function | ELU |
| Epoch | [85], [30], [4] |

***PathDSP (GEx, T) – Reactome***

| Hyperparameter | Search Grid |
|---|---|
| 1st hidden layer | [2048, 800] |
| 2nd hidden layer | [800, 512] |
| 3rd hidden layer | [512, 256] |
| 4th hidden layer | [256, 128] |
| Drop rate | [0.05, 0.1, 0.2] |
| Learning rate | [1e-5, 1e-4, 5e-4, 1e-3] |
| Weight decay | default* |

* The weight decay of the Ada Max optimizer was set to default, since the original study did not report a value for the weight decay.

| Hyperparameter | Value (LPO, LCO, LDO) |
|---|---|
| Hidden layers | [800, 800, 512, 256], [800, 800, 256, 128], [800, 800, 256, 128] |
| Learning rate | [5e-4], [1e-4], [1e-4] |
| Weight decay | default |
| Drop out | 0.1 |
| Optimization function | Ada Max |
| Activation function | ELU |
| Epoch | [89], [21], [22] |
